# Machine learning analysis identifies *Drosophila Grunge/Atrophin* as an important learning and memory gene required for memory retention and social learning

**DOI:** 10.1101/157610

**Authors:** Balint Z Kacsoh, Casey S. Greene, Giovanni Bosco

**Author notes:** Corresponding author: (GB).

## Abstract

High throughput experiments are becoming increasingly common, and scientists must balance hypothesis driven experiments with genome wide data acquisition. We sought to predict novel genes involved in Drosophila learning and long-term memory from existing public high-throughput data. We performed an analysis using PILGRM, which analyzes public gene expression compendia using machine learning. We evaluated the top prediction alongside genes involved in learning and memory in IMP, an interface for functional relationship networks. We identified *Grunge/Atrophin* (*Gug/Atro*), a transcriptional repressor, histone deacetylase, as our top candidate. We find, through multiple, distinct assays, that *Gug* has an active role as a modulator of memory retention in the fly and its function is required in the adult mushroom body. Depletion of *Gug* specifically in neurons of the adult mushroom body, after cell division and neuronal development is complete, suggests that *Gug* function is important for memory retention through regulation of neuronal activity, and not by altering neurodevelopment. Our study provides a previously uncharacterized role for *Gug* as a possible regulator of neuronal plasticity at the interface of memory retention and memory extinction.

## INTRODUCTION

A unique and fundamental characteristic of higher order organisms is the capability to recall and remember past experiences. The ability to remember past experiences is advantageous to an organism when it is presented with challenges bearing resemblance to previous experiences. The biology of memory formation and memory retention has been probed with a vast array of experimental techniques that range from behavioral testing, to physiologically correlates, to experimental perturbation through the use of chemical, anatomical, genetic, and pharmacological stresses (Greenspan 1995, 747-750). Genetic manipulations, coupled with behavioral assays, have provided important insight into memory in- and output, allowing one to understand the genetic and molecular factors that govern memory formation, retention, and retrieval. These experimental approaches provide clues to evolutionarily conserved mechanisms that could allow for vertical integration (Dubnau and Tully 1998, 407-444; Tully *et al.* 1994, 35-47).

*Drosophila melanogaster* has proven to be an important genetic model system for understanding the genetic basis of memory (Greenspan 1995, 747-750; Davis 2005, 275-302; Margulies *et al.* 2005, R700-R713; McGuire *et al.* 2005, 328-347; Davis and Zhong 2017, 490-503). The biological process of long-term memory in Drosophila has been analyzed through the use of associative and non-associative assays. Work using associative learning paradigms has provided genes and genetic functions that are now looked upon as “classical” learning. These memory genes were found through the use of associative-fear-conditioning assays (McGuire *et al.* 2005, 328-347; Tully 1987, 330-335; Tully *et al.* 1994, 35-47; Dubnau and Tully 1998, 407-444). These gene candidates were further elucidated and expanded in other assays, including the use of non-associative olfactory memory (Das *et al.* 2011, E646-54; McCann *et al.* 2011, E655-62; Ramaswami 2014, 1216-1229).

Social learning, or information exchange between a trained teacher and a naïve student, has been studied in insects, including bees (Alem *et al.* 2016, e1002564; Loukola *et al.* 2017, 833-836) and Drosophila. Studies involving food choice demonstrate that naïve, student flies can gain information through visual cues by observing trained, teacher flies, informing students that subsequently prefer the food of the trainer (Battesti *et al.* 2012, 309-313). Another form of social learning in Drosophila involves the use of the courtship-conditioning paradigm. Here, male flies that unsuccessfully court either virgin females or mated females learn to associate pheromonal cues with the courtship rejection and subsequently suppress courtship of all female flies (Siegel and Hall 1979b, 3430-3434; Ejima *et al.* 2007, 599-605; Ejima *et al.* 2005, 194-206; Siegel and Hall 1979a, 3430-3434).

Collectively, these studies in memory formation, retention, and social learning are critical in elucidating genetic and physiological mechanisms of learning and memory. However, these assays use non-ecologically relevant forms of stimuli, and thus, the genes identified might only represent a small fraction of learning and memory genes in the insect brain.

Other forms of memory have also been investigated utilizing what are now known as “classical” memory genes, which involve ecologically relevant forms of stimuli to the fly life cycle. In nature, Drosophila larvae are regularly infected by endoparasitoid wasps. In wild *D. melanogaster* populations, upwards of 90% of fly larvae are found in a wasp-infected state, suggesting they exert extremely strong selection pressures on Drosophila populations (Driessen *et al.* 1989, 409-427; Fleury *et al.* 2004, 181-194; LaSalle 1993, 197-215). Drosophila larvae can mount a cellular immune response following infection (Carton and Nappi 1997, 218-227), or adult *D. melanogaster* females can alter egg-laying (oviposition) behavior after an encounter with endoparasitoid wasps. A change in oviposition behavior entails at least two very different and quantifiable behavioral responses. First, if high ethanol containing food is made available to adult Drosophila, then female flies in the presence of wasps will actively prefer to lay eggs on ethanol-laden food (Kacsoh *et al.* 2015a, 1143-1157; Kacsoh *et al.* 2013, 947-950). This behavior persists even after the wasp threat is removed, demonstrating a memory formation and retention (Kacsoh *et al.* 2015a, 1143-1157; Kacsoh *et al.* 2013, 947-950). Second, Drosophila females depress their oviposition rate in the presence of wasps and maintain this depression following wasp removal. Wasp exposed flies will also communicate the wasp threat to naïve flies that have never seen the wasp threat, which in turn will also depress their oviposition rate, demonstrating a form of social learning (Kacsoh *et al.* 2015b, 10.7554/eLife.07423; Lefevre *et al.* 2012, 230-233; Lynch *et al.* 2016). Visual cues from the wasp or wasp-exposed teachers are sufficient to trigger these behaviors, suggesting the presence of an innate circuit yielding these behaviors (Kacsoh *et al.* 2015b, 10.7554/eLife.07423; Kacsoh *et al.* 2015a, 1143-1157; Kacsoh *et al.* 2013, 947-950). These behaviors persist in wild-type flies for multiple days after wasp encounter, allowing one to probe questions regarding memory formation and retention. The formation of these long-term memories is regulated, in part, by previously identified learning and memory genes, such as *Orb2* and *rutabaga* (Kacsoh *et al.* 2015b, 10.7554/eLife.07423; Kacsoh *et al.* 2015a, 1143-1157; Kacsoh *et al.* 2013, 947-950).

To date, the identified learning and memory genes may only represent a small fraction of the range of learning and memory genes and gene functions in the insect brain required for memory acquisition, retention, and recall. Thus, it is valuable to define and mechanistically identify novel genes and gene products. In order to facilitate a broader understanding of genes and gene products that govern memory formation, retention, and recall, the development of new methodologies are needed to identify biologically important functions that are independent of classical mutagenesis-based approached.

In the sequencing age, vast compendiums of existing and publically available data present the opportunity to address important biological questions (Faith *et al.* 2007, e8; Yan *et al.* 2010, e12139; Chikina *et al.* 2009, e1000417). Given the data abundance and availability, the question becomes, how does one utilize so much data to ask specific questions? Genome-wide expression data capture a vast array of conditions in a wide range of organisms, biological processes, and tissues. Data-mining algorithms applied to such large data sets can uncover novel, biologically relevant functions of genes that might be otherwise overlooked as candidates (Greene *et al.* 2014, 1896-1900; Greene and Troyanskaya 2012, 95-100). PILGRM (the <u>P</u>latform for Interactive <u>L</u>earning by <u>G</u>enomics <u>R</u>esults <u>M</u>ining) is a data-mining platform that allows its users to encode specific questions about a biological function of interest into user created gene sets. Probing for a specific biological function is performed by the user curating a gold standard of genes that are relevant to a particular pathway or process, which PILGRM uses for a supervised machine learning analysis of global microarray expression collections (Greene and Troyanskaya 2011, W368-74). The user can also provide a list of negative control genes, allowing for improved prediction specificity. PILGRM then trains a support vector machine (SVM (Joachims 2006, 217-226)) classifier with these gene sets to discover novel relevant genes (Greene and Troyanskaya 2011, W368-74). The SVM analysis assigns relatively high weights to conditions that differentiate gold standard (positive) genes from those in the negative standard (Greene and Troyanskaya 2011, W368-74). Thus, this machine learning approach can be used to identify novel gene targets/functions in order to further elucidate a biologically meaningful process, such as memory.

In this study, we use PILGRM to identify novel genetic candidates by using a gold standard comprised of classically-identified learning and memory genes. We examined the top prediction from PILGRM, a histone-deacetylase, *Grunge* (*gug*, CG6964), in a *Drosophila melanogaster* functional relationship network and found that it was highly connected to learning and memory genes and that the network was enriched for that particular process. We then used the natural wasp predator system to probe a role for this previously untested gene in *D. melanogaster* learning and memory by using three distinct learning and memory paradigms. These three assays probed for long-term memory formation, long-term memory retention, and social learning. Collectively, we test the hypothesis that *gug* may be a high-level regulator of learning and memory. Histone-acetylation activity has been previously identified as a regulator of mouse hippocampal learning and memory (Mews *et al.* 2017), where histone acetylation yields activation of early memory genes. Perturbation of histone acetylation activity presents defects in memory formation. A functional role for histone deacetylase activity in Drosophila memory is consistent previous reports showing that *Rpd3* (HDAC1) functions in the Drosophila mushroom body to facilitate long-term memory formation (Fitzsimons and Scott 2011, e29171), suggesting that *gug* is a strong candidate for investigation.

## RESULTS

### PILGRM ANALYSIS REVEALS NOVEL GENE TARGETS

In order to identify novel genes involved in learning and memory, we turned to the PILGRM data-mining tool (<u>http://pilgrm.princeton.edu</u>). The system requires a list of positive and negative standards. We input a positive, gold standard as well as a list of randomly selected negatives provided by the PILGRM server. Our positive standard list was comprised of previously identified learning and memory genes. Specifically, we input the genes: *for* (Donlea *et al.* 2012, 2613-2618), *Akt1* (Guo and Zhong 2006, 4004-4014), *Fmr1* (Kanellopoulos *et al.* 2012, 13111-13124; Kacsoh *et al.* 2015b, 10.7554/eLife.07423; Kacsoh *et al.* 2015a, 1143-1157), *S6k* (Vargas *et al.* 2010, 1006-1011), *CrebB-17A* (Perazzona *et al.* 2004, 8823-8828), *Adf1* (DeZazzo *et al.* 2000, 145-158; Kacsoh *et al.* 2013, 947-950; Kacsoh *et al.* 2015a, 1143-1157), *amn* (Aldrich *et al.* 2010, 33-41; Kacsoh *et al.* 2015b, 10.7554/eLife.07423; Kacsoh *et al.* 2015a, 1143-1157), *rut* (Scheunemann *et al.* 2013, 17422-17428; Kacsoh *et al.* 2015a, 1143-1157; Kacsoh *et al.* 2015b, 10.7554/eLife.07423), *Nf1* (Li *et al.* 2013, 5821-5833), *CaMKII* (Mehren and Griffith 2006, 686-689), *orb2* (Krüttner *et al.* 2012, 383-395; Kacsoh *et al.* 2015b, 10.7554/eLife.07423; Kacsoh *et al.* 2015a, 1143-1157), and *shi* (Iyengar *et al.* 2011, 883-900)(Supplementary file 1). This list of gold standards is comprised of learning and memory genes that have been validated in associative and non-associative memory assays. A subset of the genes selected here has been shown to be involved in the paradigms used in this study. We hypothesized that the use of genes that have been validated in multiple assays would provide a list of output genes most likely to provide a novel hit.

We performed the analysis utilizing PILGRM’s *Drosophila melanogaster* compendium, which consists of gene expression data sets from the Gene Expression Omnibus (GEO). Following the analysis, PILGRM provides a visual interpretation for precision of predictions at different recall thresholds (Figure 1 A). Following analysis, users are also provided a visual representation of true positive rate at various false positive rate thresholds. We find in our analysis that the area under the curve (AUC), shown as the shaded region, is 0.8130 (Figure 1 B). PILGRM also provides a list of novel predictions (Figure 1 C, Supplementary file 2). In this case, the top novel prediction is the gene *Grunge* (*gug*), whose molecular function is listed as a histone deacetylase (Wang *et al.* 2006, 525-530; Zhang *et al.* 2013, 575-583; Zhang *et al.* 2002, 45-56; Wang *et al.* 2008, 555-562; Yeung *et al.* 2017, 10.7554/eLife.23084). This gene has no previously described role in learning and memory in Drosophila. However, the Drosophila gene expression atlas (Fly:Atlas) reports *Gug* expression as most highly enriched in the fly brain (Chintapalli *et al.* 2007, 715-720). We identify other genes in our top novel predictions from PILGRM that may also have a role in memory. For example, our third target, *Twins* (*tws*), is shown to be involved in neuroblast development (Chabu and Doe 2009, 399-405). Our fourth target is TBP-associated factor 4 (Taf4), important in dendrite morphogenesis (Parrish *et al.* 2006, 820-835). Also in our top 10, we find Syndecan (Sdc), a gene involved in synapse growth (Chanana *et al.* 2009, 11984-11988). Collectively, we find our top candidates highly enriched in neurons and brain tissue with previously characterized genes having neuronal function.

**Figure 1.**
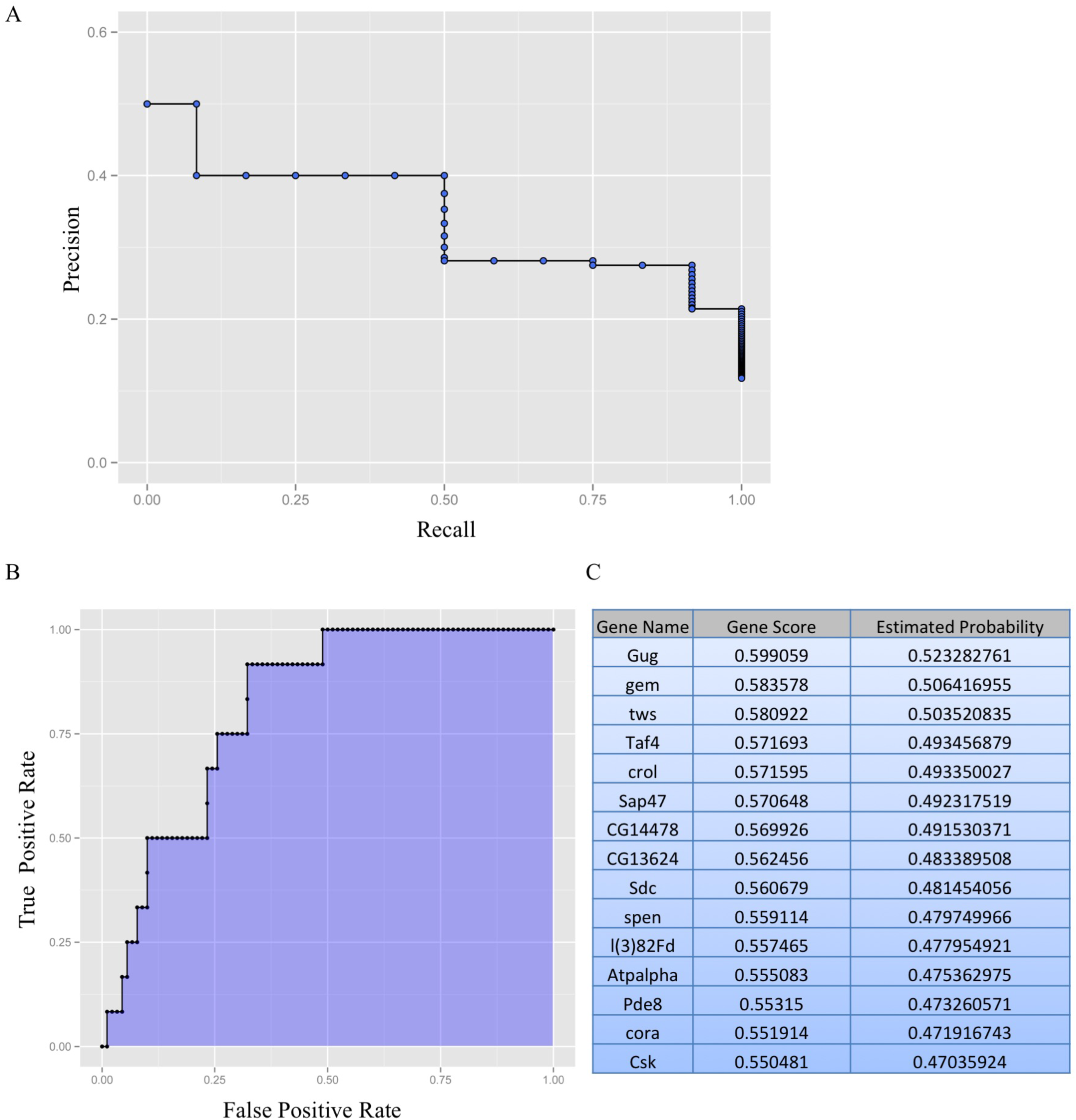
PIGLRM analysis reveals genes that may have a role in Drosophila memory, including the histone deacetylase, *Gug*. Using known learning memory genes as the positive standard, we performed PILGRM analysis to identify a list of novel target genes that may be involved in memory. The analysis provides visual analysis of precision of predictions at different recall thresholds (A), visual analysis of true positive rate at various false positive rate thresholds (the area under the curve, shown as the shaded region, for this analysis is 0.8130) (B), and the top novel predictions based on the analysis (C).

We became intrigued with the possibility of histone modifications playing a role in fly memory. Recent work has shown that aberrations in mouse hippocampal histone acetylation yields mice with impaired long-term spatial memory formation, a cognitive process that relies on histone acetylation to activate early memory genes (Mews *et al.* 2017; Kandel *et al.* 2014, 163-186; Korzus *et al.* 2004, 961-972; Gräff and Tsai 2013, 97-111; Wood *et al.* 2005, 111-119). In Drosophila, the HDAC1 homolog, *Rpd3*, also is implicated in learning and memory functions (Fitzsimons and Scott 2011, e29171). Given these observations, in conjunction with observations showing epigenetic deregulation as a mechanism for neuropsychiatric diseases (Kandel *et al.* 2014, 163-186; Zovkic *et al.* 2013, 61-74; Gräff and Tsai 2013, 97-111; Walker *et al.* 2015, 112-121), we wished to elucidate the possibility of *Gug* playing a role in learning and long-term memory.

### *GUG* SHOWS HIGH CONNECTIVITY TO KNOWN LEARNING AND MEMORY GENES

In order to further examine *Gug* before moving to genetic experiments, we used the integrative <u>m</u>ulti-species <u>p</u>rediction (IMP) webserver to determine the extent to which *Gug* is connected with our known learning and memory gene set (Wong *et al.* 2012, W484-90; Wong *et al.* 2015, W128-33). IMP integrates gene-pathway annotations from the selected organism in addition to mapping functional analogs. This system has been shown to provide accurate gene-process predictions (Chikina and Troyanskaya 2011, e1001074).

We queried the network with *Gug* and our gold standard set. We set the network filter to only show interactions whose minimum relationship confidence was 0.85, a stringent threshold. We find that *Gug* is connected, both directly and indirectly, to our known learning and memory genes (Figure 2). Unsurprisingly, this network is enriched for the biological processes of long-term memory (26.3%, p-value: 1.29e-8), short-term memory (15.8%, p-value: 1.85e-6), learning (21.1%, p-value: 1.21e-5), and associative learning (18.4%, p-value: 4.51e-5). Given the PILGRM and IMP results, we sought to test the role of *Gug* in a variety of assays probing for its role in memory formation, retention, and social learning.

**Figure 2.**
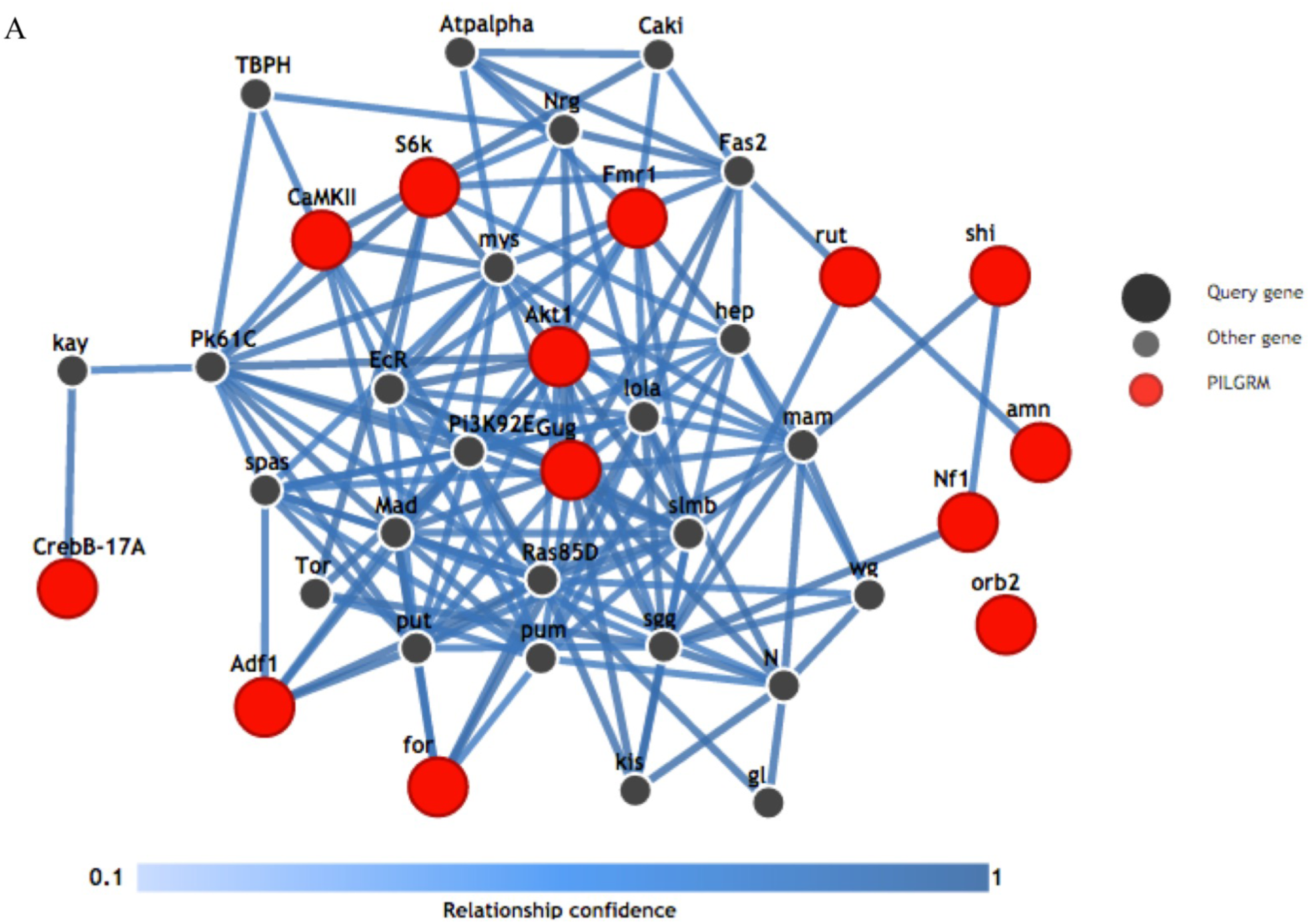
IMP analysis indicates high network connectivity of *Gug* and known memory genes. A visual representation of a gene network utilizing our positive standards from Figure 1 with gug shows a high degree of network connectivity. Lines indicate a minimum relationship confidence of 0.85.

### *GUG* HAS A ROLE IN LONG TERM MEMORY RETENTION

Given our bioinformatics results, we hypothesized that *gug* has a role in long-term memory formation and retention. We tested this question using two distinct assays, both involving the predatory wasp *Leptopilina heterotoma* (strain Lh14). Both behavioral assays utilize adult *D. melanogaster*, who can alter egg-laying behavior when encountering predatory wasps.

We first used an ethanol seeking food preference assay where, following a wasp exposure, female flies will actively prefer to lay eggs on ethanol-laden food (Kacsoh *et al.* 2013, 947-950; Kacsoh *et al.* 2015a, 1143-1157). Briefly, we exposed 3-5 day old flies in 3 batches of 100 female flies and 20 male flies with 50 female wasps. Control flies underwent the same process, but lacked wasps. Following a 24-hour exposure, flies and wasps were separated. Exposed and unexposed flies are transferred in batches of 5 female and 1 male flies to a fly corral, a Petri dish with holes, where the center of the two holes were 6 cm apart, and the edge of the two holes were 7.2 cm apart. We place Falcon tube caps to the holes. The caps contain 0.375 grams of flaky instant blue hydrated fly food with either distilled-water or distilled water with ethanol (6% by volume). Caps are changed every 24-hours for 3-days and egg counts were performed on caps. All counts are blinded (see methods) (Kacsoh *et al.* 2015a, 1143-1157).

We find that wild-type *D. melanogaster* continue to oviposit on ethanol laden food across each of the 3, 24-hour time points tested following wasp exposure (Figure 3 A) (Kacsoh *et al.* 2015a, 1143-1157; Kacsoh *et al.* 2013, 947-950). Interestingly, we find that *Gug* heterozygous mutants (Spradling *et al.* 1999, 135-177) (*Gug*/+) show a defect in persistence of oviposition on ethanol-laden food (Figure 3 B). These flies show a defect in memory retention, starting on day 2. Day 1 of the assay does not show a memory defect, suggesting that memory formation is not affected. By day 3, we observe no difference in exposed and unexposed flies in the mutants, but still have a strong difference in wild-type flies, suggesting memory extinction has been achieved in the heterozygotes (Supplementary Figure 2). However, these experiments cannot exclude the possibility that the *Gug* gene product is required in non-neural tissues, and that the observed behavior is not specific to neuronal function.

**Figure 3.**
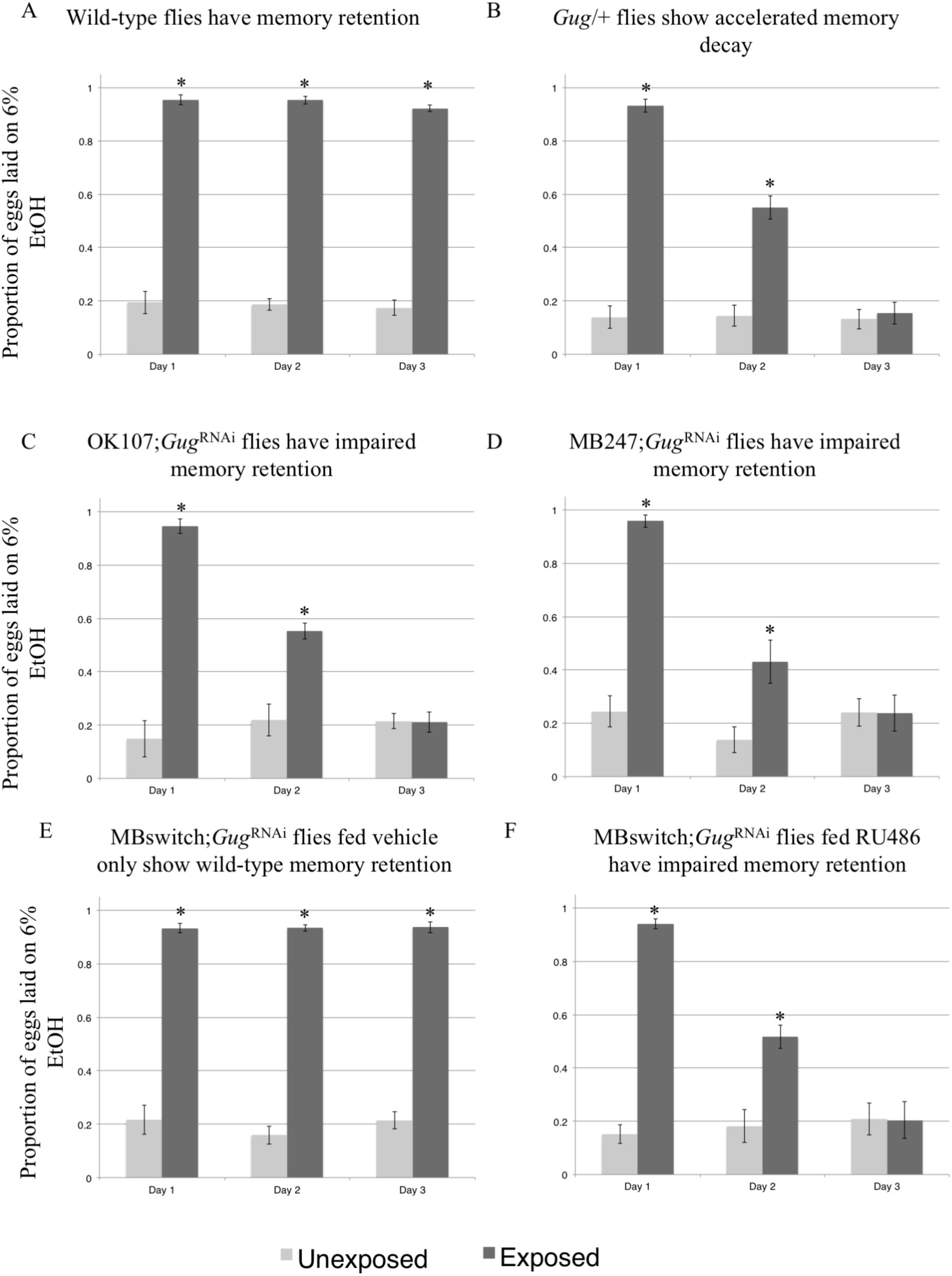
*Gug* is involved in memory retention using ethanol choice memory assay. Proportion of eggs laid on 6% ethanol oviposition cap following wasp exposure after three 24-hour time points is shown. Wild-type flies continue to oviposit on 6% ethanol after wasp exposure (A). *Gug*/+ heterozygotes (B) and flies expressing a *Gug*^RNAi^ hairpin in the MB show impaired memory retention (C-D). MB switch flies fed vehicle control show wild-type memory (E), while flies expressing a *Gug*^RNAi^ via RU486 feeding show impaired memory retention (F). Error bars represent 95% confidence intervals (n = 10 biological replicates) (*p < 0.05).

The mushroom body (MB) of the adult brain is thought to be required for behaviors that are dependent on learning and memory (Aso *et al.* 2009, 156-172; Claridge-Chang *et al.* 2009, 405-415; Schwaerzel *et al.* 2003, 10495-10502; Masse *et al.* 2009, R700-R713). The behaviors we set out to test *Gug*’s role in have been shown to be MB dependent, demonstrating a bona fide memory paradigm (Kacsoh *et al.* 2015b, 10.7554/eLife.07423; Kacsoh *et al.* 2015a, 1143-1157). A critical question arising after testing the *Gug* heterozygotes was whether *Gug* is required in the MB for memory formation and/or retention. We used the GAL4/UAS system to drive expression of an RNA-hairpin targeting *Gug* mRNA (*Gug*^RNAi^) in conjunction with either the MB driver OK107 or MB247. We find that driver lines alone and the line containing the *Gug* hairpin alone have wild-type memory retention (Supplementary Figure 1 A-C). However, flies expressing the *Gug*^RNAi^ in the MB phenocopy the *Gug* heterozygote phenotype (Figure 3 C-D), where we observe no defect in memory formation, only a defect in memory retention beginning on the second day of observation. Again, we observe memory ablation in these flies by day 3, suggesting a memory retention defect (Supplementary Figure 2).

While the MB drivers in conjunction with the *Gug*^RNAi^ presented a memory retention phenotype, the constitutive expression of the hairpin presents the possibility of *Gug*’s function being essential for early neuronal development. This suggestion means that the knockdown phenotypes may simply reflect developmental defects that preclude proper adult MB functions. In order to test elucidate this possibility, we turned to the GAL4-based Gene-Switch system where the GAL4 transcription factor is fused to the human progesterone ligand-binding domain (Burcin *et al.* 1999, 355-360). We used flies expressing the Gene-Switch transgene specifically in the MB where only upon an administration of the pharmacological Gene-Switch ligand, RU486, GAL4 transcription factor would become active (Mao *et al.* 2004, 198-203). We find that feeding of RU486 to the outcrossed MB GeneSwitch driver line yields a wild-type memory retention phenotype. This demonstrates that feeding of the GeneSwitch ligand does not perturb memory formation or retention (Supplementary Figure 1 D). We also observe wild-type memory retention when the MB GeneSwitch is in combination with the *Gug*^RNAi^ when fed vehicle control (methanol) (Figure 3 E). When fed the RU486 solution, these flies express the *Gug*^*RNAi*^ hairpin and show the same memory retention defect as when the *Gug*^RNAi^ transgene is in conjunction with constitutively active MB driver, and the *Gug/+* heterozygote (Figure 3 E). Again, these flies show no memory formation defect when compared to wild-type. However, there is accelerated memory decay at day 2 and the memory is ablated by day 3, demonstrating a defect in memory retention (Supplementary Figure 2). Thus, we believe that the memory retention defect observed is a result of *Gug* being a necessary gene product in the MB to facilitate memory retention in the adult Drosophila brain. Given these data, we use the MB GeneSwitch line to drive expression of the *Gug* hairpin for all subsequent experiments. This approach allows us to delineate the role of *Gug* in memory retention from other important functions it may also perform during development.

We decided validate our conclusion indicative of the role of *Gug* in memory retention by utilizing a second, distinct memory assay. This second assay utilizes another *D. melanogaster* behavior following wasp exposure, where Drosophila female flies depress their oviposition rate following wasp exposure. Flies are given only standard Drosophila media as a substrate, with no food choice in this assay. The oviposition depression following wasp exposure is observed for multiple days after the predator threat is removed (Kacsoh *et al.* 2015b, 10.7554/eLife.07423; Lefevre *et al.* 2012, 230-233). We utilized Fly Condos to measure oviposition rate (Kacsoh *et al.* 2015b, 10.7554/eLife.07423). Briefly, *D. melanogaster* were exposed for 24 hours to wasps in cylindrical 7.5cm long by 1.5cm diameter tubes of the Fly Condos (Genesse). Each tube contains five female flies and one male fly, either with three female wasps (exposed) or with no wasps at all (unexposed) (see methods). After 24-hours, wasps are removed, and flies are placed into new, clean Fly Condos. We repeat this transfer for each of 3 days following wasp exposure. At every 24-hour time point, food-plates are removed, replaced with new food plates, and embryos are counted in a blinded manner.

Consistent with previous observations, wild-type flies depress oviposition in the presence of wasps and following wasp removal (Figure 4 A, Supplementary Figure 2 B) (Lefevre *et al.* 2012, 230-233; Kacsoh *et al.* 2015b, 10.7554/eLife.07423). We find that *Gug* heterozygotes (*Gug*/+) have wild-type memory on day 1 following wasp exposure, but show accelerated memory decay, where by day 2 there is no difference between exposed and unexposed groups (Figure 4 B, Supplementary Figure 4). In order to ascertain the specificity of the phenotype with respect to memory, we again utilized the MB GeneSwitch line. We find that the *Gug*^RNAi^ construct outcrossed to *Canton S* has wild-type memory retention (Supplementary Figure 3 A) (Kacsoh *et al.* 2015b, 10.7554/eLife.07423). When the *Gug*^RNAi^ is expressed in conjunction with the MB GeneSwitch fed vehicle only, we observe wild-type memory retention (Figure 4 C). When this line is fed RU486, leading to *Gug* knock-down, we observe a memory retention defect comparable to the heterozygote, where day one is unaffected, suggesting that memory formation is wild-type, but day two begins to show accelerated memory decay, where again we observe no difference between exposed and unexposed (Figure 4 D, Supplementary Figure 4). These data show no defect in memory formation, only memory retention. Given the data from the food choice and egg retention assays, we observe that *Gug* has a functional role in the MB, independent of memory formation, and modulates memory retention.

**Figure 4.**
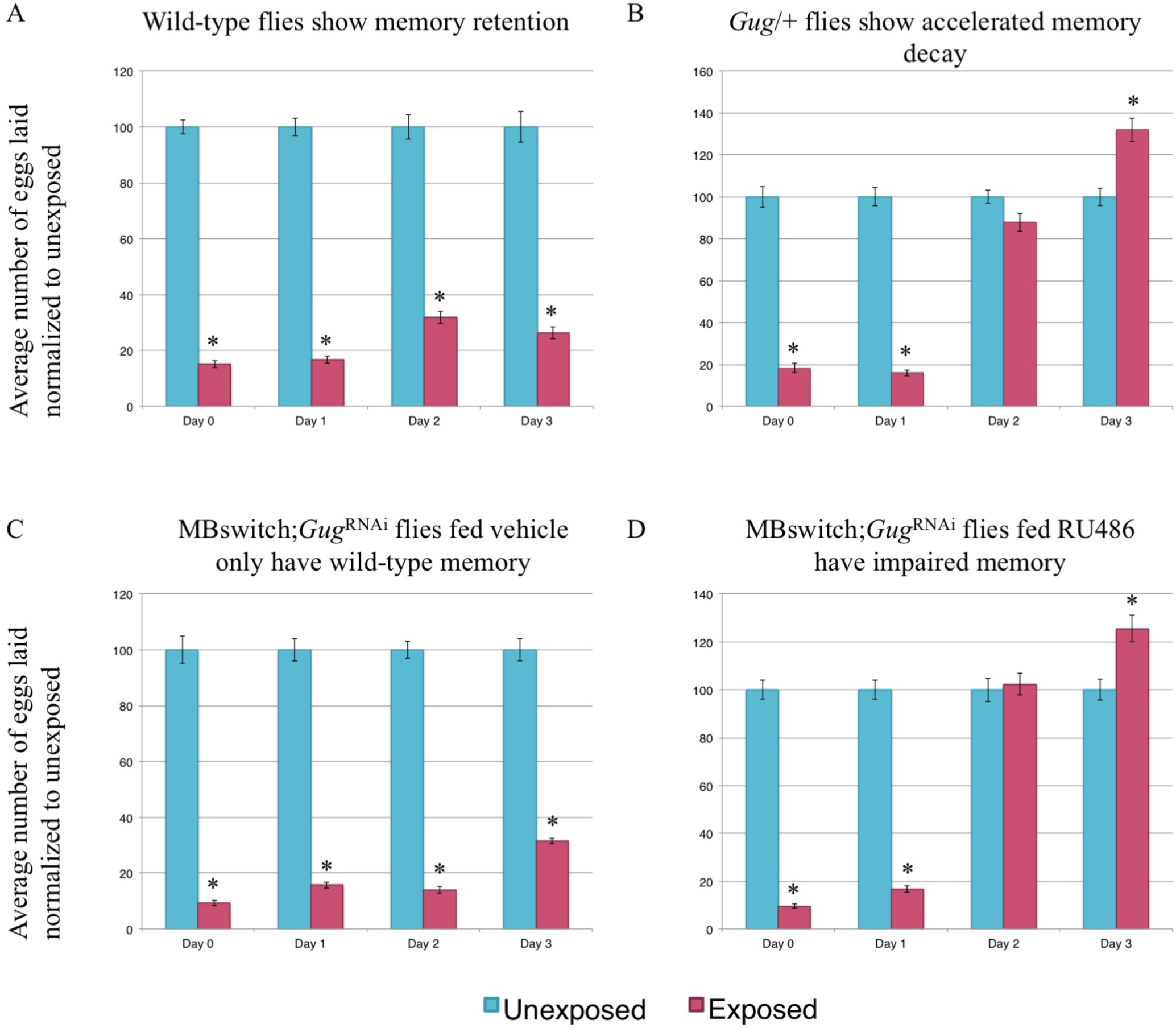
*Gug* is involved in memory retention using egg retention assay. Percentage of eggs laid by exposed flies normalized to eggs laid by unexposed flies is shown. Wild-type flies exposed to wasps lay fewer eggs than unexposed flies for multiple days (A). *Gug*/+ flies have impaired memory retention (B). MB switch flies fed vehicle control show wild- type memory (C), while flies expressing a *Gug*^RNAi^ via RU486 feeding show impaired memory retention (D). Error bars represent standard error (n = 24 biological replicates) (*p < 0.05).

### *GUG* HAS A ROLE IN TEACHING BEHAVIOR

We observe a memory retention, but not formation, defect in *Gug* heterozygous mutants and *Gug* knockdown in the MB. We sought to ask whether this memory retention defect translates to other behaviors that are present following wasp exposure, including a social learning phenomenon. Following wasp exposure, flies depress oviposition and communicate the threat of wasps to naïve, unexposed flies. We term the experienced flies as “teachers” and the naïve flies “students.” We hypothesized that the memory retention defects observed (Figures 3-4, Supplementary Figures 2 and 4) may translate to defects in teaching behavior. To ask this, we utilized the Fly Condos (Kacsoh *et al.* 2015b, 10.7554/eLife.07423). Briefly, we exposed wild-type flies to wasps for 24-hours, followed by wasp removal. We then placed the exposed and unexposed teacher flies into new Fly Condos with three naïve female flies expressing Histone-RFP (His-RFP) for an additional 24-hours (see methods). Following a 24-hour co-incubation, we removed His-RFP students, placed exposed and unexposed teachers into new Fly Condos containing a new batch of 3 female naïve His-RFP flies. We repeated this process 3 times following wasp exposure as a means of testing the maintenance and ability of teaching behavior. New batches of students were used at each 24-hour time point. At every 24-hour time point, food-plates were removed, replaced with new food plates, and embryos were counted in a blinded manner. The His-RFP line was ideal for discriminating mixed populations of non-RFP and RFP embryos, allowing us to probe specificity of student and teacher behavior.

Consistent with previous data, we find that wild-type flies can instruct multiple batches of students across the 3 days tested (Figure 5 A) (Kacsoh *et al.* 2015b, 10.7554/eLife.07423). We find that *Gug* heterozygotes (*Gug*/+) have wild-type teaching ability on day 1, which quickly decays on days 2 and 3 (Figure 5 B). To again probe for neuronal specificity, we turned to the MB GeneSwitch line. We find that outcrossed *Gug*^RNAi^ flies have wild-type teaching behavior across each time-point tested (Supplementary Figure 5). When the *Gug*^RNAi^ is expressed in conjunction with the MB GeneSwitch line fed vehicle only, we observe wild-type teaching behavior across all 3 days (Figure 5 C). When this line is fed RU486, allowing for expression of *Gug*^RNAi^ transgene, we find wild-type teaching ability on day 1, but find decaying teaching ability across days 2 and 3, similar to the heterozygote (Figure 5 D, Supplementary Figure 6). Collectively, the data suggests that memory of the wasp exposure must be maintained in order to exhibit persistent teaching behavior. *Gug* is not required for teaching behavior on day 1, where the flies still depress oviposition. Once the memory begins to its erasure, we observe the accelerated decay in teaching behavior. Interestingly, in *Gug* deficient flies, we observe no difference between exposed and unexposed oviposition on day 2. However, these flies are still able to teach, albeit not as efficiently as wild-type. By day 3, *Gug* deficient flies are unable to teach naïve students, while wild-type flies are still efficient teachers (Figure 5, Supplementary Figure 6). The data suggest that there may be two circuits in the fly brain governing 1) wasp memory and 2) teaching behavior, both of which require *Gug* to maintain signaling.

**Figure 5.**
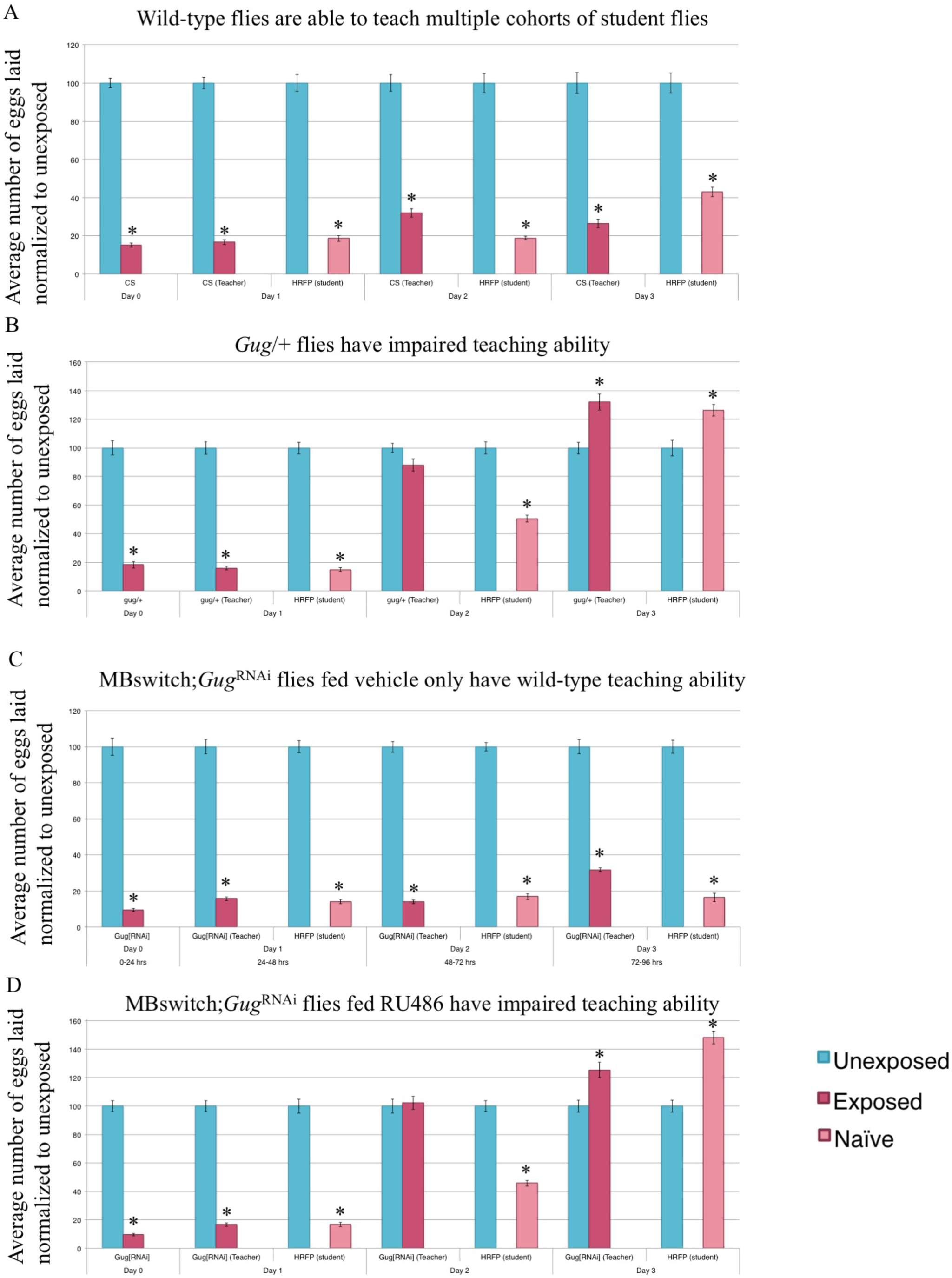
*Gug* is involved in teaching ability. Percentage of eggs laid by exposed flies normalized to eggs laid by unexposed flies is shown. Wild-type flies exposed to wasps can teach multiple student cohorts (RFP-Histone) across three days (A). *Gug*/+ flies have impaired teaching ability (B). MB switch flies fed vehicle control show wild-type teaching ability (C), while flies expressing a *Gug*^RNAi^ via RU486 feeding show impaired teaching ability (D). Error bars represent standard error (n = 24 biological replicates) (*p < 0.05).

### *GUG* HAS A ROLE IN STUDENT BEHAVIOR

We observe a memory retention phenotype when flies lack *Gug* in the MB. We observe no defect in memory formation. Given these results, we wished to ask whether a different type of learning and memory paradigm is also affected following *Gug* removal. To do this, we utilized the social learning paradigm described above, but used *Gug* deficient flies as students to see how they learn from wild-type teachers. Since both *Gug* heterozygotes and RNAi depleted Drosophila were able to perceive and respond to wasp presence and form the initial memory, we hypothesized that *Gug* deficient flies may also be able to learn from wasp-exposed teacher flies. An alternative hypothesis is that learning in the context of social interactions with other flies could be different from learning that takes place directly from wasp-exposure. In this latter case, the molecular, cellular and/or neuronal requirements for social learning might be different than those for non-social learning. We propose this given that the inputs, a wasp or a teacher fly, are very different for the observer fly given the differences in the visual, olfactory, and other understudied cues being detected.

In order to test the role of *Gug* in social learning, we modified the Fly Condo oviposition depression assay. We exposed His-RFP flies to wasps for 24-hours, followed by wasp removal. We then placed the exposed and unexposed teacher flies into new Fly Condos with three naïve female flies that were either wild-type, or *Gug* deficient, for an additional 24-hours. Following a 24-hour co-incubation, food-plates were removed, and embryos were counted in a blinded manner.

Consistent with previous results, we find that wild-type flies are able to learn from His-RFP teacher flies (Figure 6 A) (Kacsoh *et al.* 2015b, 10.7554/eLife.07423). Interestingly, we find that *Gug* heterozygous (*Gug*/+) students have an impaired social learning ability when compared to wild-type students (Figure 6 B). These flies are still able to learn from teachers, but at a much less efficient rate when compared to wild-type students. We wished to test the role of *Gug* in the MB in a social learning context. We find that the outcrossed *Gug*^RNAi^ show wild-type learning ability (Supplementary Figure 7). When expressed in combination with the MB GeneSwitch line and fed vehicle only, we observe wild-type learning (Figure 6 C). When this line is fed RU486, inducing *Gug*^RNAi^ transgene expression, we observe an impaired social learning ability, similar to the heterozygote (Figure 6 D). We note that the ability to learn from a teacher is not ablated in *Gug* deficient flies, but instead impaired (Supplementary Figure 8). In other assays used, the day 1 phenotype has always been similar to wild-type, but in this assay, there is a distinct difference in the day 1 phenotype of these deficient flies. Collectively, these data strongly suggest that *Gug* has an important function in social learning and information processing. Again, this protein function occurs specifically in the MB information-processing center of the fly brain. The data also suggest that the type of learning taking place from wasp exposure may be fundamentally different from the type of learning taking place during social learning.

**Figure 6.**
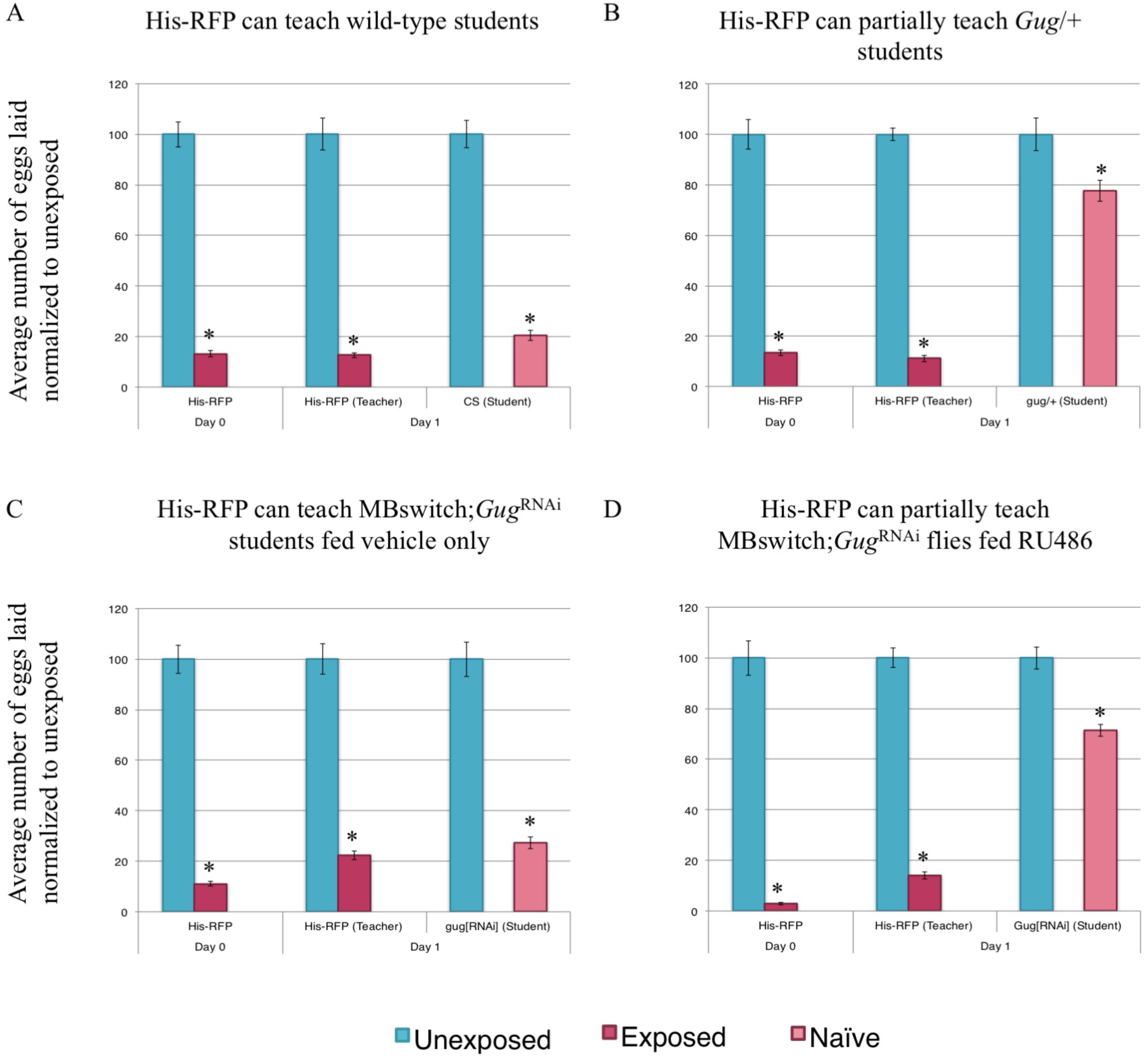
Gug has a role in social learning. Percentage of eggs laid by exposed flies normalized to eggs laid by unexposed flies is shown. Wild-type flies (His-RFP) exposed to wasps can teach wild-type (CS) students (A). *Gug*/+ flies have impaired learning ability (B). MB switch flies fed vehicle control show wild-type learning ability (C), while flies expressing a *Gug*^RNAi^ via RU486 feeding show impaired learning ability (D). Error bars represent standard error (n = 24 biological replicates) (*p < 0.05).

## DISCUSSION

In this study, we have presented the use of a machine-learning platform, PILGRM, in concert with a functional network analysis tool, IMP, to identify and evaluate a novel regulator of learning and memory in Drosophila. We identified *Gug*, a transcriptional repressor known to complex with the Rpd3/HDAC histone deacetylase (Fitzsimons and Scott 2011, e29171). To our knowledge, *Gug* itself has not been previously implicated to function in *Drosophila* memory.

We tested *Gug*’s role in memory formation and memory retention through two unique assays. Through these assays we find wild-type memory formation following training. However, memory retention is severely perturbed when the protein is missing in the fly mushroom body. We note that this function in memory retention for *Gug* is different from what has previously been reported for Rpd3, and therefore constitutes a novel function for *Gug* specifically in maintenance of long-term memory, rather than formation. Our observation raises the interesting possibility that the HDAC activity of Rpd3 recruited to chromatin by Gug could also be important in memory retention. It is important to note that the memory defects in *Gug* deficient flies we observe are not simply a result of developmental defects, given the data utilizing the MB GeneSwitch lines. In these lines, the depletion of *Gug* in adult MB neurons takes place after cell division and development has been completed. These data suggest that *Gug* has an active role in post-mitotic neurons throughout memory retention and/or retrieval in the fly MB, most likely via its transcriptional repressor function.

We highlight that memory formation is not affected in flies lacking *Gug*, as the day 0 and day 1 data looks like wild-type. This also lack of difference also suggests that the sensory cues required to identify a wasp as a threat, visual or otherwise, are intact in these flies lacking *Gug*. Thus, these flies are able to process the relevant input information from the wasp signal to form a memory. Thus, *Gug* is not required to detect the input or to make the initial memory. This result also suggests that the wiring and development of the fly brain remains intact to produce a functioning brain, given that the initial signal and interpretation are wild-type. We observe phenotypic differences on days 2 and 3, where we find the rapid extinction of memory, or accelerated memory decay. Wild-type flies show strong memory retention following the exposure, but in each of the assays we use, we find total memory ablation by day 3.

Additionally, we find that perturbations in *Gug* yield differences in social behavior. We find that teaching behavior in *Gug* deficient flies that is normal on day one when compared to wild-type. As the memory decays in these flies, so does the teaching behavior, suggesting that the retention of both memory and teaching behavior is modulated by *Gug* activity. Of greatest interest, we find that social learning is also perturbed in *Gug* deficient flies, where students are not able to effectively gain information from wild-type teacher flies. These flies still have an ability to function as a student, but this ability is severely impaired. This observation is important because it suggests that molecular requirements for learning in a social context from experienced flies may require fundamentally different molecular and/or neuronal circuitry, and it is therefore distinct from learning in non-social contexts. Interestingly, humans with trinucleotide expansions within the *Atrophin 1* gene, the human *Gug* homolog, exhibit autism-like behaviors (Licht and Lynch 2002, 51-54). Thus, the observation of a social learning deficiency in flies could serve as a model for studying autism.

Recent work has shown that visual cues are sufficient to elicit a wasp-response and a teacher-student response (Kacsoh *et al.* 2013, 947-950; Kacsoh *et al.* 2015b, 10.7554/eLife.07423). In this study, for each assay used, we provided direct interaction between insects (either fly-wasp or teacher-student), where visual, olfactory, tactile, and auditory cues may be exchanged. Thus, we cannot rule-out the possibility that *Gug* may have an effect on certain sensory components of the nervous system. It remains possible that a chromatin transcriptional repressor may alter the excitability of the neurons involved in each of these circuits to make the neuron less excitable. A less excitable neuron may lead to a less responsive fly, where wild-type flies achieve maximal excitation and form a stable memory, while *Gug* deficient flies are trained to a lesser degree resulting in an unstable memory that decays faster. This lowered neuron excitation may also account for the student-teacher interaction, where the *Gug* deficient flies do not learn as effectively as wild-type due to sensory dampening. This alternative hypothesis does not affect the conclusion that *Gug* is required in the MB for memory retention, only that *Gug* might also function elsewhere in the learning circuitry. Future experiments that utilize isolated sensory stimuli would be able to examine these possibilities.

Previous work using a murine hippocampal model demonstrates a role for histone acetylation and subsequent expression of neuronal genes involved in memory formation (Mews *et al.* 2017). In mammals, Sp3 has been shown to recruit HDAC2 to synaptic specific genes, without interfering in other HDAC2 gene targets (Yamakawa *et al.* 2017, 1319-1334). Together, these findings suggest the presence of specific recruitment factors that dictate the acetylation state of cell-type specific genes. In Drosophila, *Gug* (*Atrophin*) may act in a similar way by recruiting HDAC1/2 in neurons specifically (Licht and Lynch 2002, 51-54). We propose a model where histone acetylation and deacetylation are acting in concert to promote neuronal plasticity (Figure 7). Given our results, we hypothesize that deacetylase activity is important in the memory retrieval process, where certain gene activity is turned off, or repressed. This repression may help elongate the memory retention, as without deacetlyase activity, we observe accelerated memory decay. The targets of *Gug* in the MB may serve as memory extinction genes (Abel and Lattal 2001, 180-187; Rudenko *et al.* 2013, 1109-1122), or genes that promote memory decay, and repression of these genes promotes memory maintenance. Our data also suggest that histone deacetylase activity is dispensable for memory formation, suggesting that other mechanisms govern the initial input acquisition.

**Figure 7.**
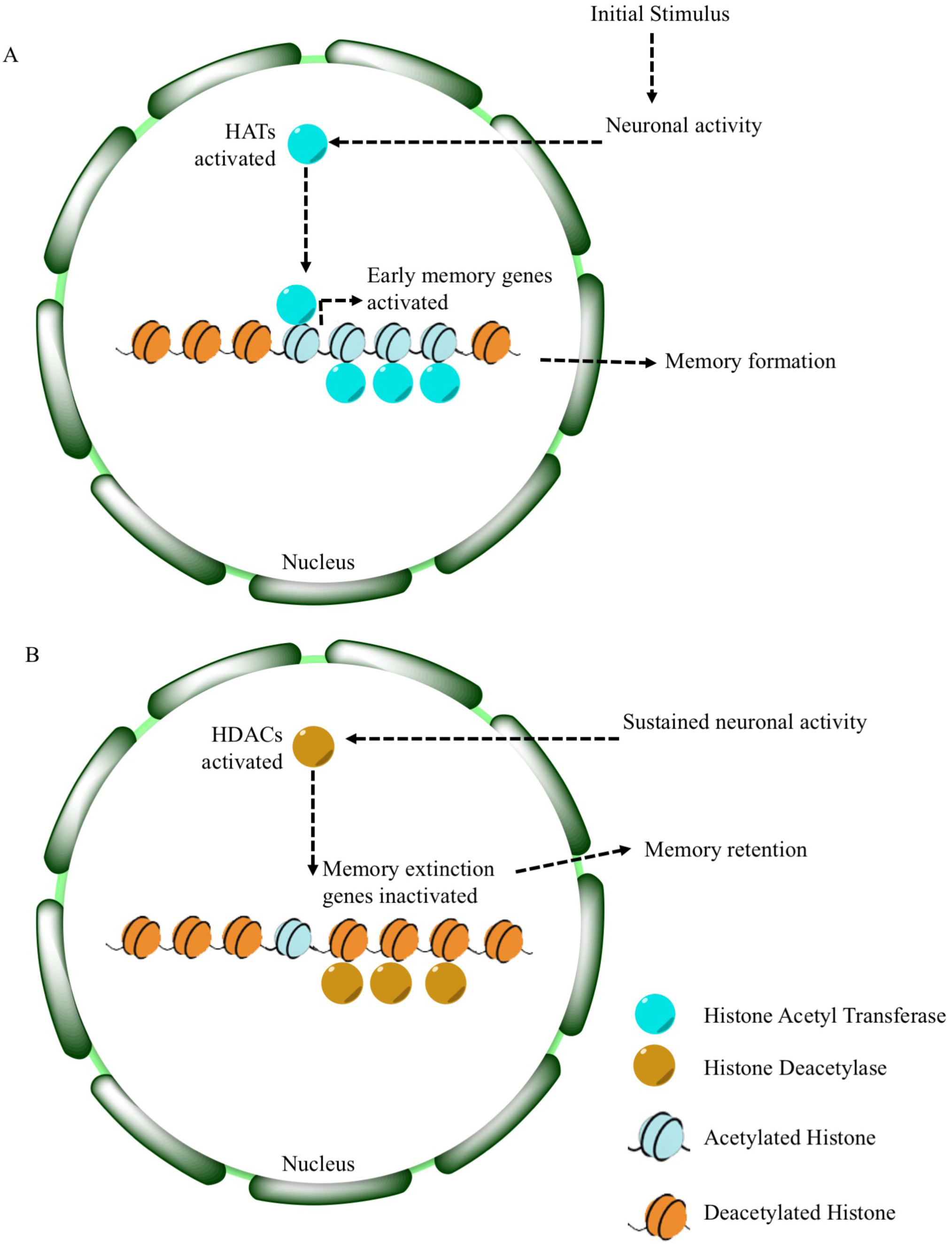
Proposed model for histone deacetylation in memory retention. Model for learning and memory shows a role for *Gug*. At the start of memory formation, histone acetylases are upregulated following neuronal stimulation, promoting upregulation of transcription of memory related genes (A). Following memory consolidation, these gene sites on chromatin may become deacetylated, in addition to memory extinction genes, to promote memory retention (B).

Collectively, our data highlights the value of machine-learning approaches for biological function prediction via biologist-driven analyses. PILGRM enables data-driven experimentation in conjunction with knowledge-based discovery through the selection of a biologically relevant gold standard. Given that PILGRM utilizes a large collection of public gene expression data, it may identify genes that are not differentially regulated in an investigator’s own experiment, but act in unison with curated genes of interest. It can also identify consistent patterns across a compendium. This approach may help to identify regulators where small, potentially non-statistically significant changes in transcript abundances can have biologically meaningful changes. These changes can be subtle, yet could lead to neuron plasticity. Epigenetic mechanisms are continually found to be important regulators of neuronal plasticity and functional output. These mechanisms are implicated in maintaining neuronal homeostasis, and perturbations have been implicated in neuropsychiatric diseases (Walker *et al.* 2015, 112-121; Mews *et al.* 2017; Gräff and Tsai 2013, 97-111; Kandel *et al.* 2014, 163-186). Our study provides a previously uncharacterized role for *Gug* as a possible regulator of neuronal plasticity at the intersection of memory retention and memory extinction.

## Acknowledgements

We thank Greg Roman, FlyBase, and the Bloomington Drosophila Stock Center, for stocks. We thank Sassan Hodge for technical assistance. We thank Julianna Bozler for technical and graphical assistance. We thank two anonymous reviewers for helpful comments on the manuscript. We acknowledge grants from Geisel School of Medicine at Dartmouth, a Gordon and Betty Moore Foundation Data-Driven Discovery Investigator grant GBMF4552 (CSG), the National Institute of Health Pioneer grant 1DP1MH110234 (GB), the National Science Foundation DBI-145830 (CSG and GB), and the Defense Advanced Research Projects Agency grant HR0011-15-1-0002 (GB).

## SUPPLEMENTARY FIGURE LEGENDS

**Supplementary Figure 1.**
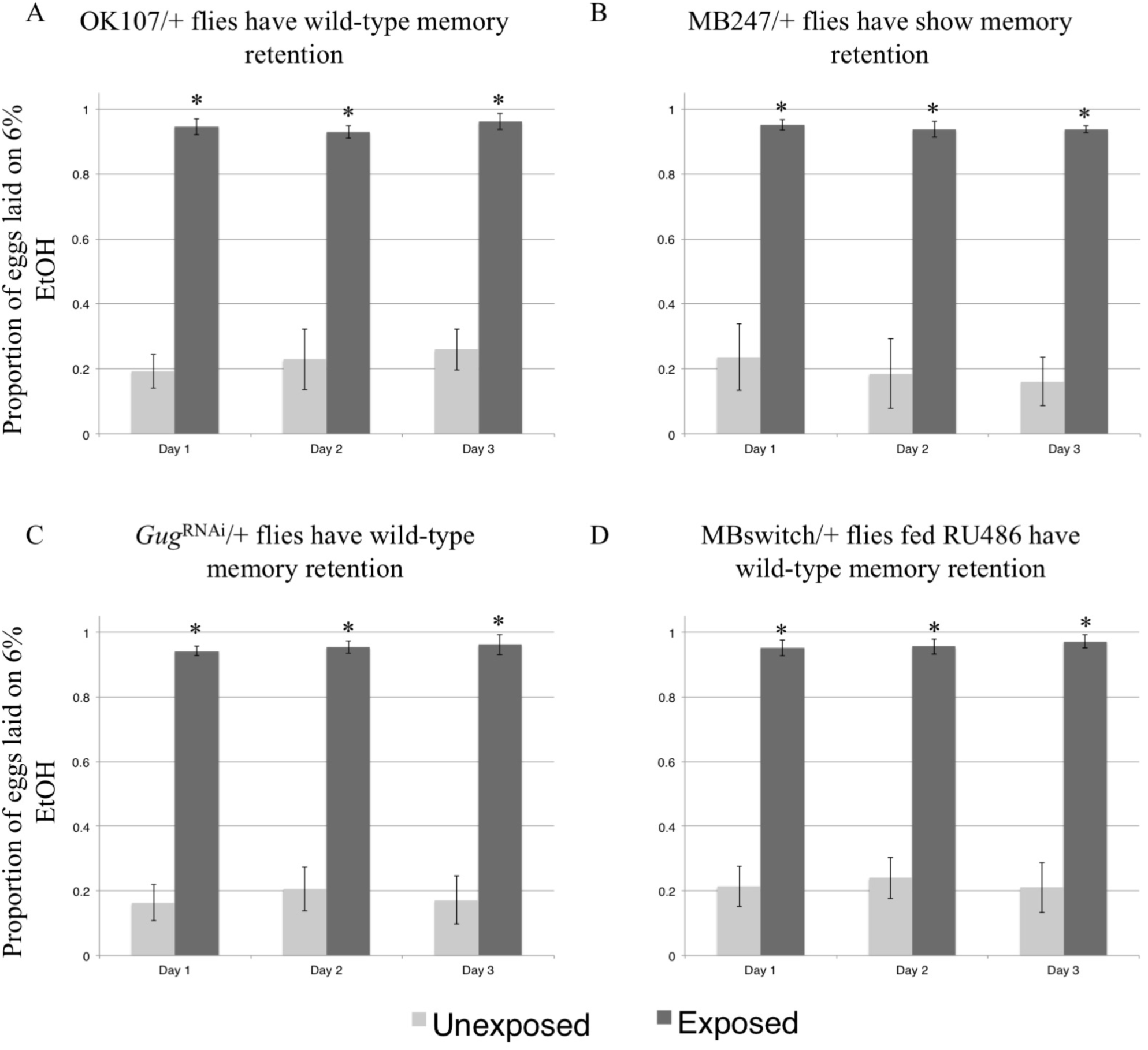
Further evidence demonstrating the role of *Gug* in memory retention. Proportion of eggs laid on 6% ethanol oviposition cap following wasp exposure after three 24-hour time points is shown. Controls outcrossed to Canton S continue to oviposit on 6% ethanol after wasp exposure. Outcrossed genotypes shown are OK107/+ (A), MB247/+ (B), *Gug*^RNAi^ (C), and MBswitch/+ fed RU486. Error bars represent 95% confidence intervals (n = 10 biological replicates) (*p < 0.05).

**Supplementary Figure 2.**
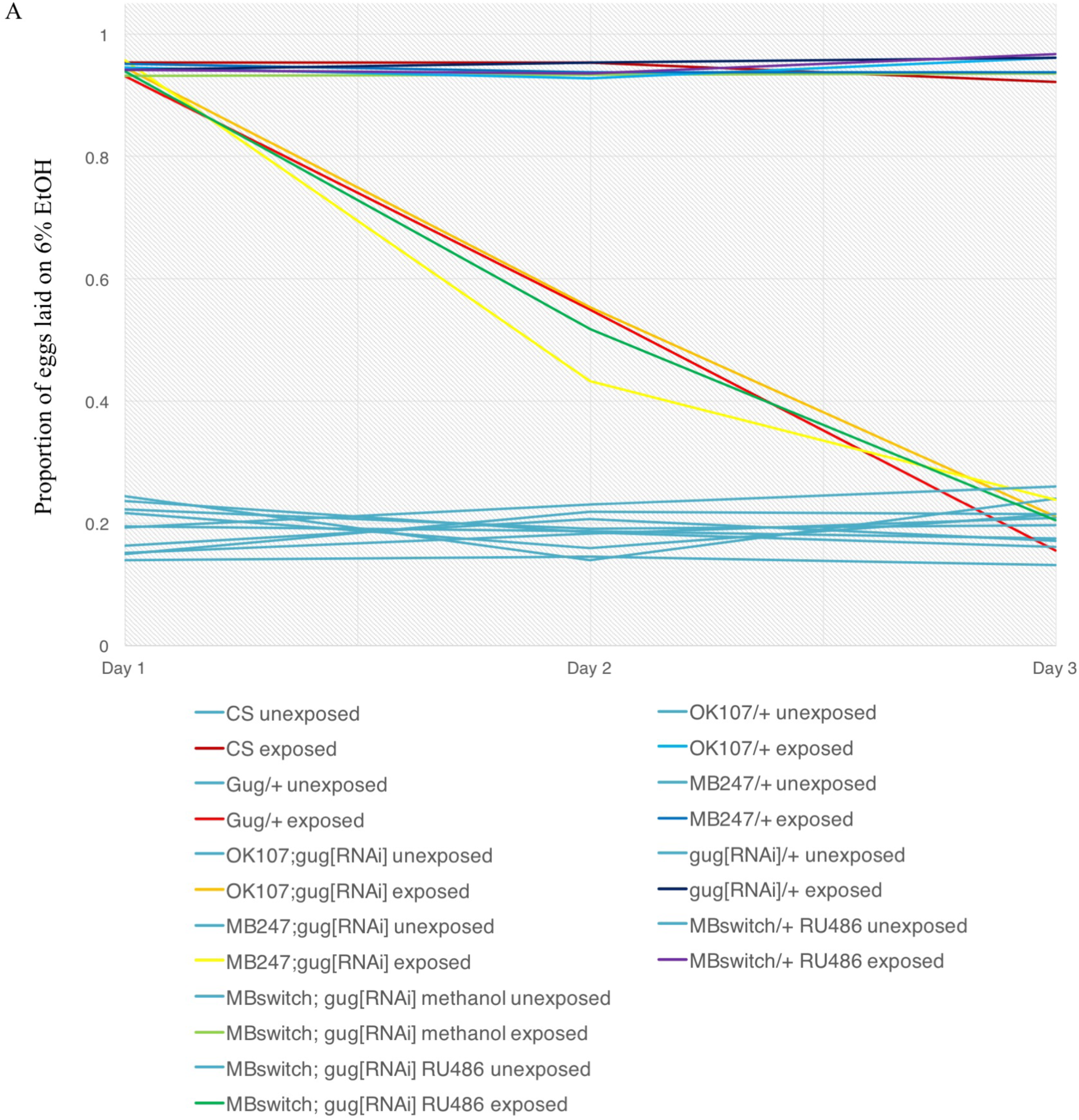
Collective evidence demonstrating the role of *Gug* in memory retention. Proportion of eggs laid on 6% ethanol oviposition cap following wasp exposure after three 24-hour time points is shown utilizing line graphs. These data are presented as bar graphs in Figure 3 and Supplementary Figure 1. All control lines are shown in light blue, while all exposed treatments are shown in varying colors. *Gug* perturbation leads to an accelerated memory decay with the curves going down to unexposed levels.

**Supplementary Figure 3.**
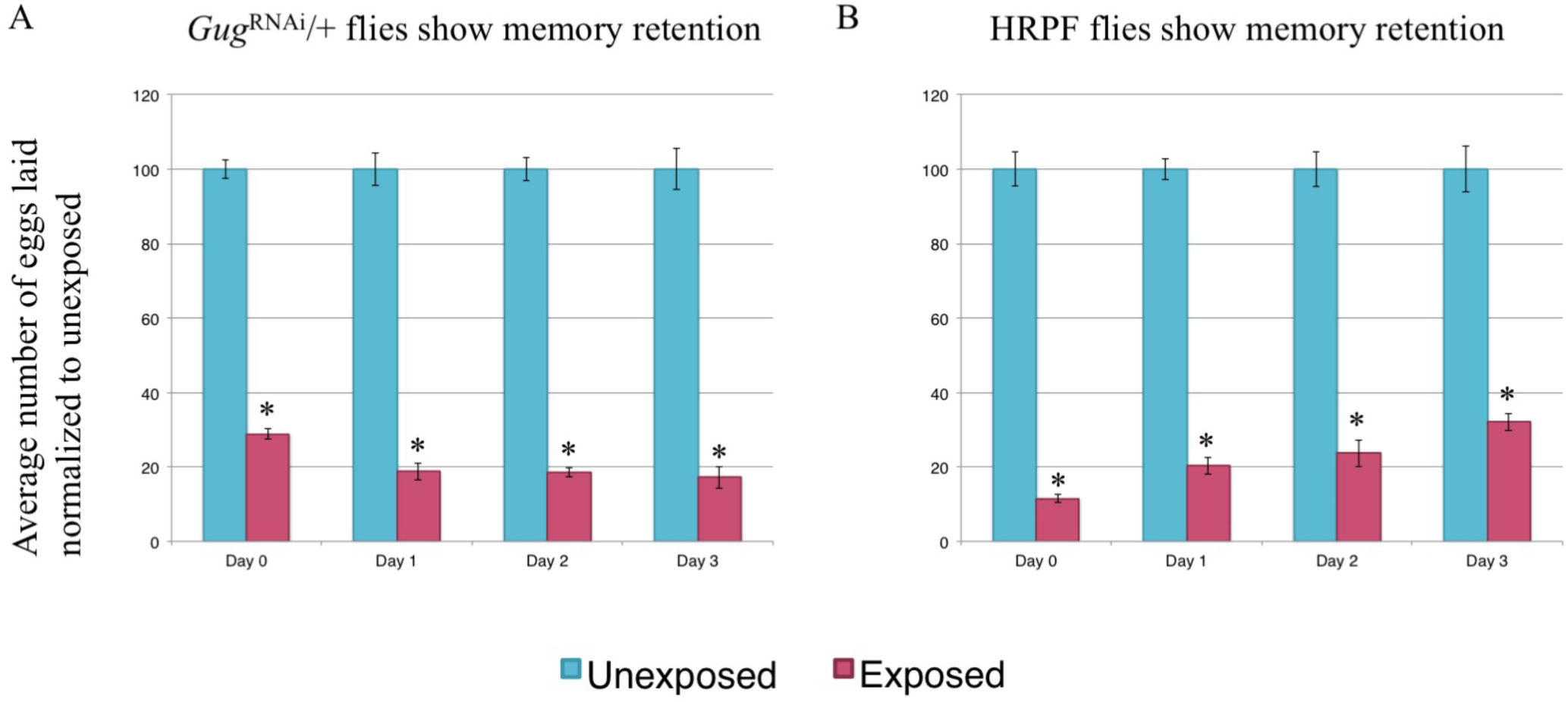
Further evidence demonstrating the role of *Gug* in egg-retention memory. Percentage of eggs laid by exposed flies normalized to eggs laid by unexposed flies is shown. Wild-type flies exposed to wasps lay fewer eggs than unexposed flies for multiple days (A). *Gug*^RNAi^/+ (A) and His-RFP flies have wild-type memory retention (B). Error bars represent standard error (n = 24 biological replicates) (*p < 0.05).

**Supplementary Figure 4.**
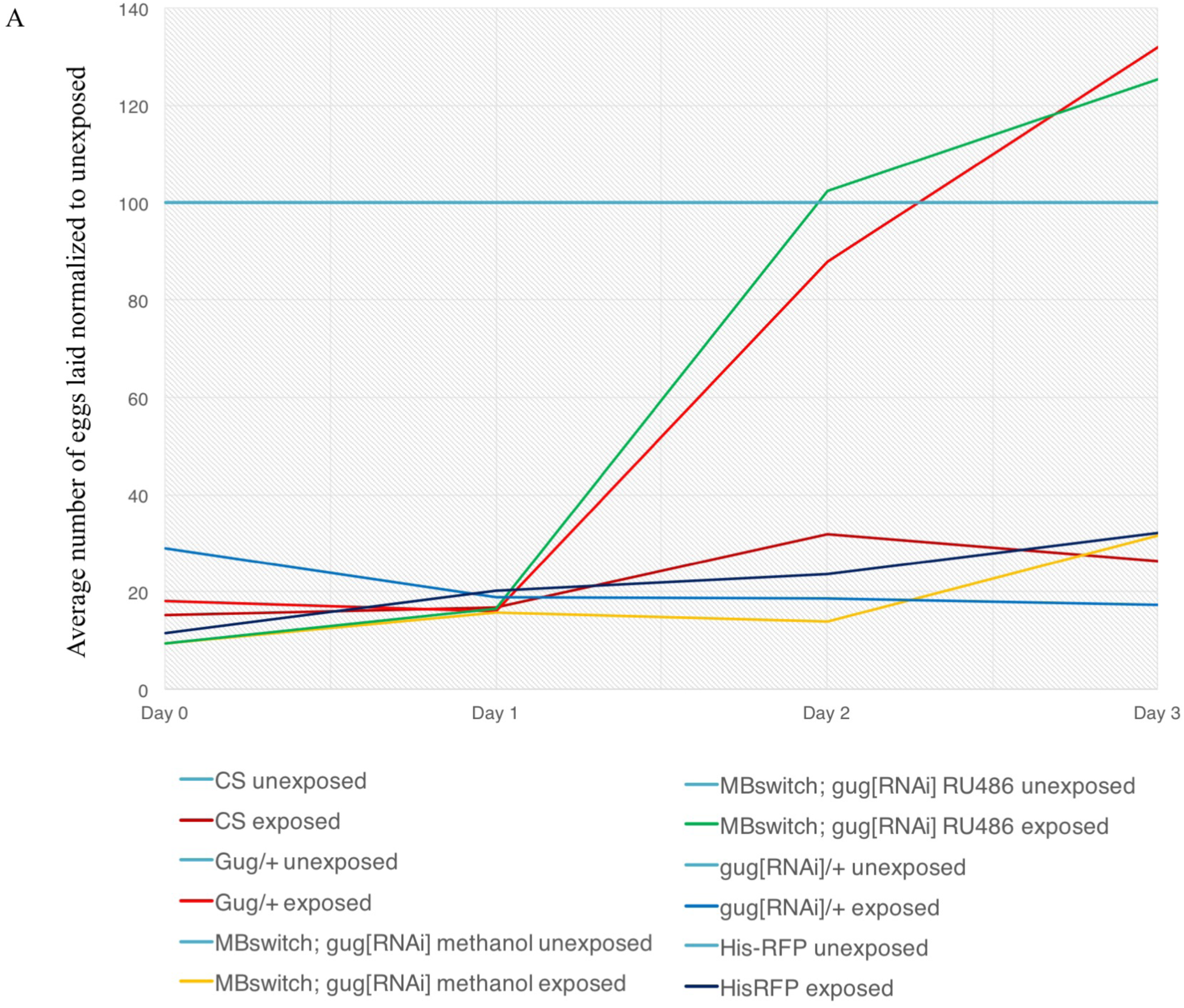
Collective evidence demonstrating the role of *Gug* in egg-retention memory. Percentage of eggs laid by exposed flies normalized to eggs laid by unexposed flies is shown utilizing line graphs. The data are also presented in bar graph form in Figure 4 and Supplementary Figure 3. All control treatments are shown in light blue, while exposed treatments are shown in varying colors. *Gug* perturbation leads to an accelerated memory decay, with the curves going up to unexposed levels.

**Supplementary Figure 5.**
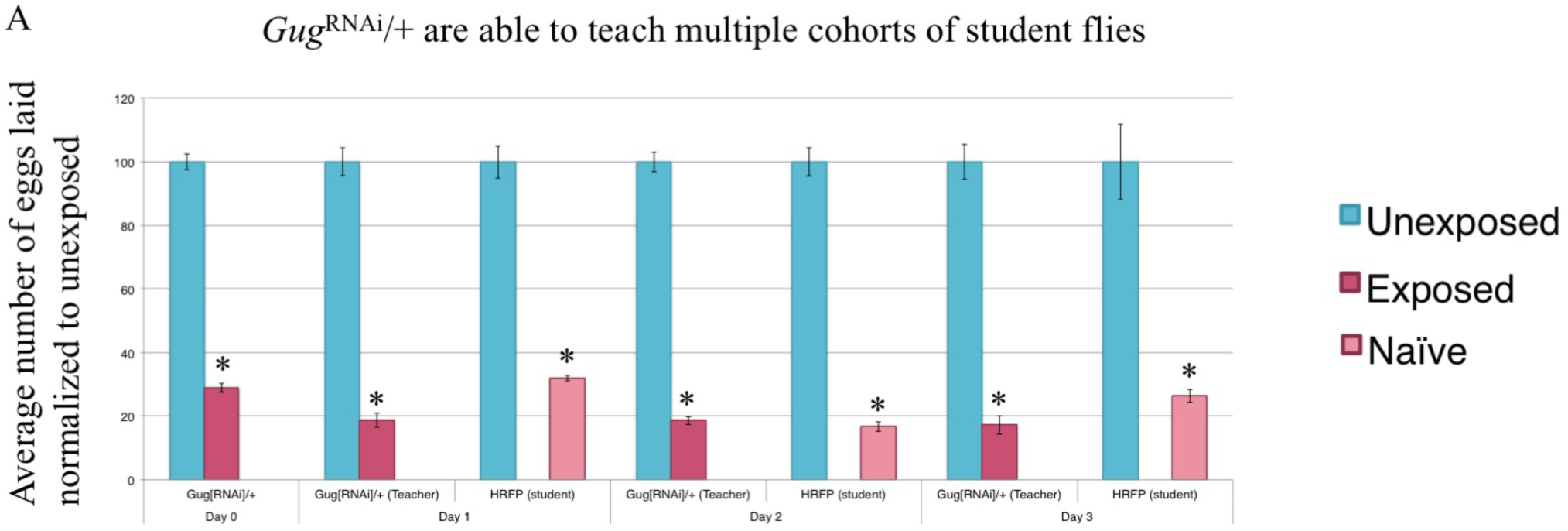
Further evidence demonstrating the role of *Gug* in teaching behavior. Percentage of eggs laid by exposed flies normalized to eggs laid by unexposed flies is shown. *Gug*^RNAi^/+ (outcrossed to Canton S) flies exposed to wasps can teach multiple student cohorts (RFP-Histone) across three days (A). Error bars represent standard error (n = 24 biological replicates) (*p < 0.05).

**Supplementary Figure 6.**
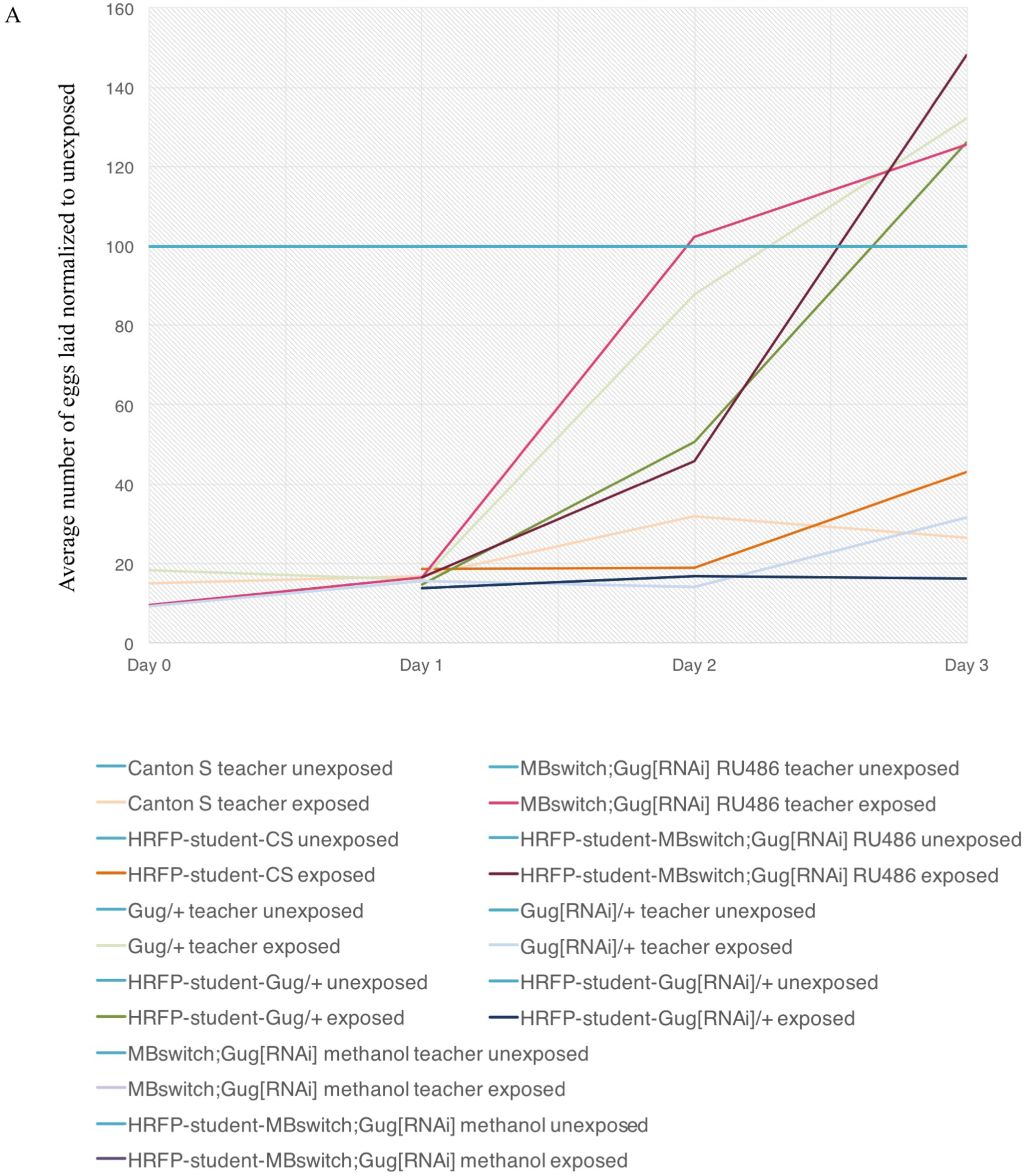
Collective evidence demonstrating the role of *Gug* in teaching behavior. Percentage of eggs laid by exposed flies normalized to eggs laid by unexposed flies is shown utilizing line graphs. The data are also presented in bar graph form in Figure 5 and Supplementary Figure 4. All control treatments are shown in light blue, while exposed treatments are shown in varying colors. A paired teacher-student is shown in the same color, with the teacher being lighter, and the student being darker (see legend). *Gug* perturbation leads to an accelerated memory decay and a loss of teaching efficiency, with both curves going up to unexposed levels.

**Supplementary Figure 7.**
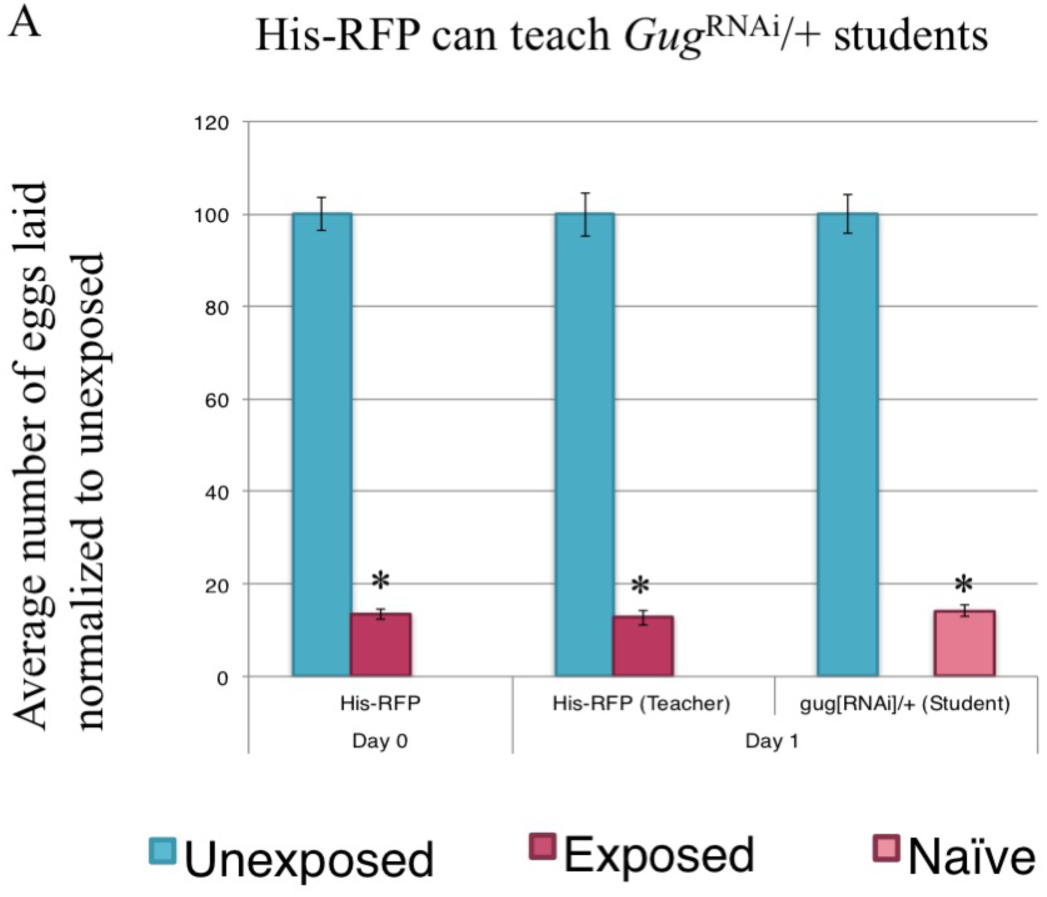
Further evidence demonstrating the role of *Gug* in social learning. Percentage of eggs laid by exposed flies normalized to eggs laid by unexposed flies is shown. Wild-type flies (His-RFP) exposed to wasps can teach *Gug*^RNAi^/+ students (A). Error bars represent standard error (n = 24 biological replicates) (*p < 0.05).

**Supplementary Figure 8.**
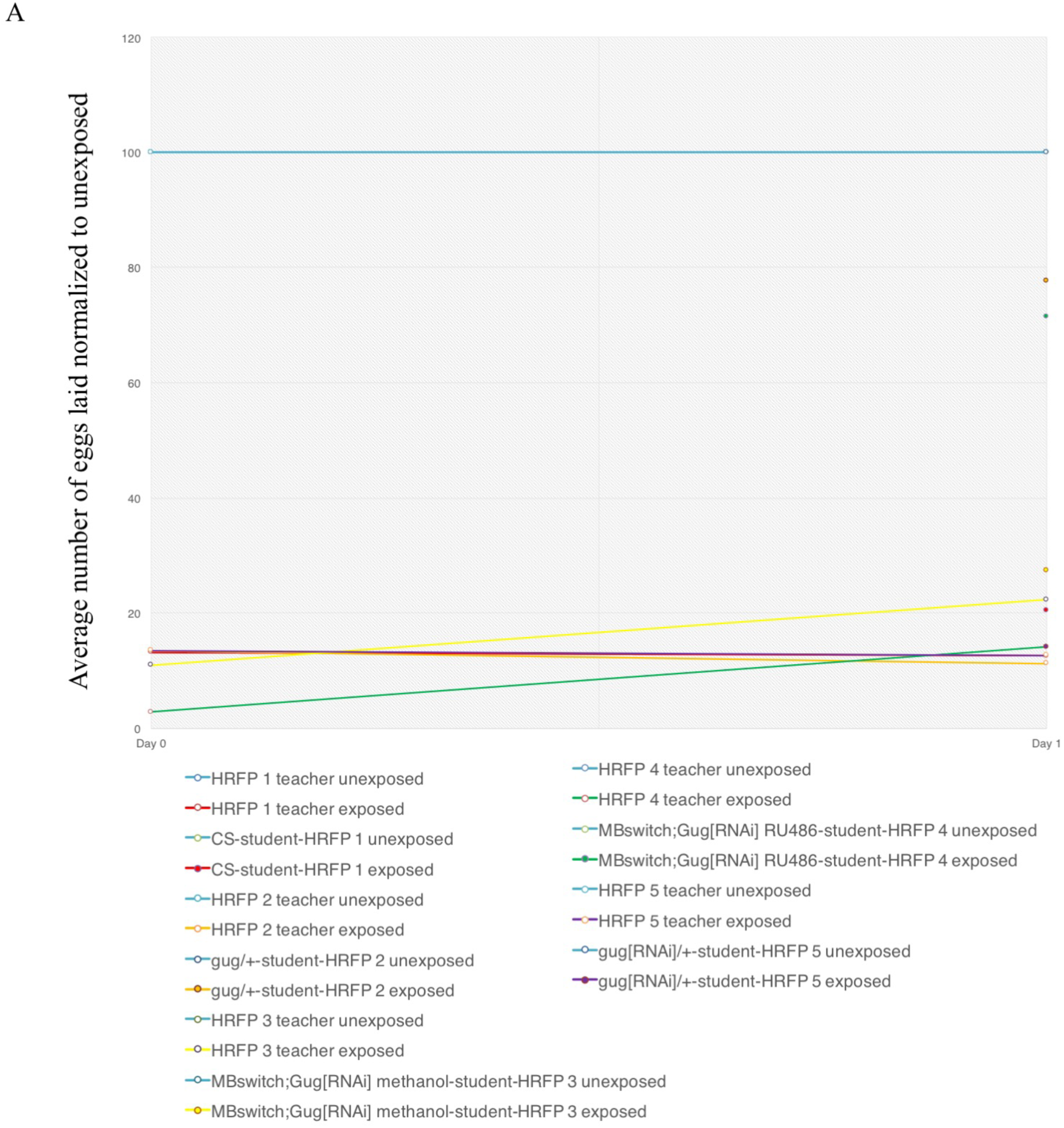
Collective evidence demonstrating the role of *Gug* in social learning. Percentage of eggs laid by exposed flies normalized to eggs laid by unexposed flies is shown in line graph form. The data are also presented in bar graph form in Figure 6 and Supplementary Figure 7. All control treatments are shown in light blue, while exposed treatments are shown in varying colors. A paired teacher-student is shown in the same color, with the teacher being lighter, and the student being darker having a filled in circle (see legend). *Gug* perturbation leads to impaired social learning.

## SUPPLEMENTARY TABLES

**Supplementary Table 1.**
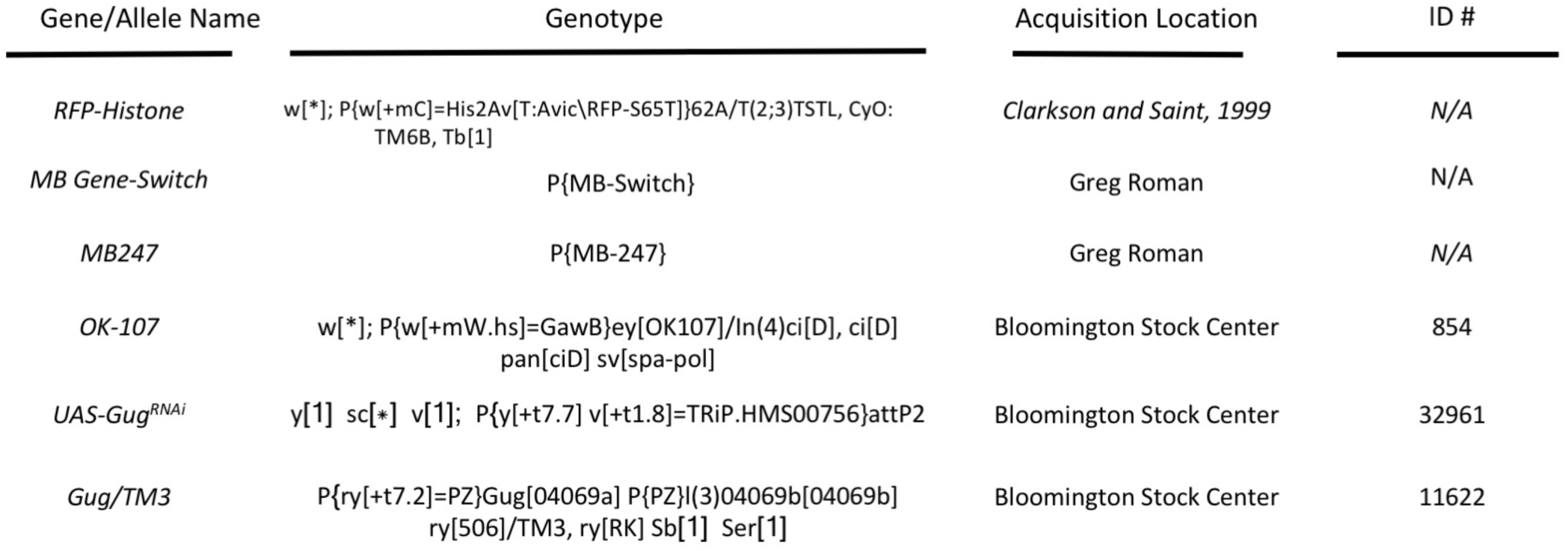
Fly genotypes used in this study. Name, genotype, acquisition location, and stock identification number (if applicable) are shown.

## SUPPLEMENTARY FILES

**Supplementary File 1. Gene list of positive and negative standards used in PILGRM analysis.**

Genes in positive standard are indicated with a (1). Genes in negative standard indicated by a (- 1).

**Supplementary File 2. Novel genes as predicted by PILGRM.**

Output gene list from PILGRM analysis.

## MATERIALS AND METHODS

### Insect Species/Strains

The *D. melanogaster* strains Canton-S (CS), and a Histone-RFP transgenic line, was used as the wild-type strain for oviposition preference after wasp exposure. The mushroom body Gene- Switch line (MB GeneSwitch) and the MB247 were kindly provided by Greg Roman (Baylor College of Medicine). Flies were maintained at room temperature with approximately 30% humidity. UAS-Gug^RNAi^, Gug/TM3 (Spradling *et al.* 1999, 135-177), and OK-107 were acquired from the Bloomington Drosophila Stock Center (stock numbers 32961, 11622, and 854, respectively) (Supplementary Table 1). Flies aged 3-5 days were used for all experiments.

All species and strains used were maintained in fly bottles (Genesse catalog number 32-130) containing 50 mL of standard Drosophila media. Bottles were supplemented with 3 Kimwipes rolled together and placed into the center of the food.

The Figitid larval endoparasitoid *Leptopilina heterotoma* (strain Lh14 (Schlenke *et al.* 2007, e158)) was used in all experiments. In order to culture wasps, batches of 14 female and 5 male adult flies (*Drosophila melanogaster*, strain Canton S) were allowed to lay eggs in standard Drosophila vials containing standard Drosophila medium supplemented with activated yeast for 5 days. After 5 days, flies were removed and replaced with adult wasps (15 female, 6 male), which then attacked the developing fly larvae. Wasp vials were supplemented with approximately 500 μL of a 50% honey/water solution applied to the inside of the vial plugs. Wasps aged 3-7 days post-eclosion were used for all experiments.

### PILGRM/IMP analysis

For the PILGRM analysis, we utilized the PILGRM version one data-mining algorithm (<u>http://pilgrm.princeton.edu</u>) (Greene and Troyanskaya 2011, W368-74). The data set used in the analysis was derived from the Fruit Fly Compendium collected in August 2011. This data was collected from the Gene Expression Omnibus. The compendium used contained a total of 3,139 arrays from 186 different experiments. This data covers 12,864 Entrez gene identifiers. Within experiments, arrays were processed using the affy (Gautier *et al.* 2004, 307-315)bioconductor (Gentleman *et al.* 2004, R80) package. The gene-expression values were then normalized and summarized using the medianpolish method (Irizarry *et al.* 2003, 102-119). The resulting experimental collections were then combined for learning using C++ Sleipnir library (Huttenhower *et al.* 2008, 1559-1561).

Our gold standard was curated to include known learning and long-term memory genes. Our negative standards were selected by PILGRM in a randomized manner. Following the identification of gug, we input our curated gold standard and gug into IMP (<u>http://imp.princeton.edu</u>) (Wong *et al.* 2012, W484-90; Wong *et al.* 2015, W128-33) with a stringent minimum gene connection of 0.85 confidence.

### Fly Oviposition Ethanol Choice

For fly memory assays, we used modified Petri dishes, termed Fly Corrals, as previously used and described (Kacsoh *et al.* 2015a, 1143-1157). Holes are drilled into the Petri dish, where the center of the two holes is 6 cm apart. The diameter of each hole is ∼1.2 cm. A nitex nylon mesh is melted onto the top of the dish in order to allow for ventilation. The mesh used has 120μm openings (Genesee Scientific catalog number 57-102). Dishes were cleaned using the Fisher Brand Sparkleen powder (catalog number 04-320-4) using a 10% Sparkleen solution.

Dishes are allowed to soak in the cleaning solution for at least 2 hours and subsequently rinsed in distilled water. Plates are then allowed to air dry for 24 hours. This cleaning protocol is followed both before and after an experiment was performed (Kacsoh *et al.* 2015a, 1143-1157).

For preparation of the oviposition media in the Fly Corrals, we measured approximately 0.375 grams of instant blue Drosophila medium (Fisher Scientific Catalog number S22315C) into the caps of 15 mL Falcon Tubes (S22315C, Biological Resource Center, No.:22315C). For control (0% ethanol) food, we pipette 2250 μL of distilled water onto the instant food. For 6% ethanol food, we pipette 1966.5 μL of distilled water onto the instant food. Subsequent to this, we pipette 141.75 μL of 95% ethanol (190 Proof (95%), USP/NF/FCC/EP/BP/JP) onto the food. After mixing the liquid and the food, we immediately place one cap containing ethanol and one cap containing no ethanol onto the cage with lab tape (VWR) (Kacsoh *et al.* 2015a, 1143-1157).

For memory assays utilizing the fly corral, 50 female flies and 10 male flies were co-incubated with 50 female Lh14 wasps for 24 hours in 2.25 cm diameter vials or sham exposed (control). Flies and wasps are then separated and flies are placed into fly corrals after the 24-hour exposure period with 5 females and 1 male fly placed per dish. Following a 24-hour period, caps are removed and replaced with freshly prepared caps (prepared in the exact manner the original caps are made). This process is repeated for 3 days following wasp exposure. Once caps are removed, the number of eggs on both the ethanol cap and the control cap are counted. Ten replicates are performed. All egg counts are blinded such that the counter is unaware of experimental condition and genotype (Kacsoh *et al.* 2015a, 1143-1157).

### Fly Oviposition Depression

We measured fly oviposition rates were using The Fly Condo™ (Genesee Scientific Cat # 59-110), as previously described (Kacsoh *et al.* 2015b, 10.7554/eLife.07423). The Fly Condo contains 24 independent chambers, where each chamber is 7.5cm long by 1.5cm diameter. Each condo has a 24-well food plate into which we dispensed 2mL of standard Drosophila media. In order to assay egg retention of flies in the presence of wasps (acute exposure), we place 5 female flies and 1 male fly into one chamber of the Fly Condo. For exposed units, 3 female Lh14 wasps are also placed in the units. The oviposition plate from control and experimental condos are counted 24 hours later. To assay memory of wasp exposure, after the 24-hour exposure, we remove all wasps and transfer all flies to a new Fly Condo. The oviposition food plate is replaced with freshly poured food (2mL). We repeat this process for three days following the wasp exposure and egg counts are performed every 24-hours. All egg counts are blinded such that the counter is unaware of experimental condition.

Social communication and social learning was tested as previously described (Kacsoh *et al.* 2015b, 10.7554/eLife.07423). In order to assay fly-fly communication and the social learning period, 5 female flies and 1 male fly are placed into one chamber of The Fly Condo in the control, while 3 female Lh14 wasps are placed with the flies in the experimental setting for 24 hours. After the 24-hour wasp or sham (control) exposure, wasps are removed and replaced with 3 female “student” flies. These are naïve flies, never having seen a wasp. Flies are placed into new, clean fly condos for the second 24-hour period. For experiments with His-RFP teachers, the experiment terminates after one batch of students. For experiments using His-RFP as students, we replace the student flies with new batches of 3 female naïve His-RFP flies. This is repeated for 3 24-hour periods following the wasp exposure. This experiment measures teaching ability of flies. The oviposition plates contain 2mL of standard Drosophila media, replaced every 24-hours.

Fly embryo counts from each plate are made at each 24-hour time points. All egg counts are blinded such that the counter is unaware of experimental condition and genotype.

All treatments are run at 25°C, at 40% humidity, with a 12:12 light:dark cycle in twenty-four replicates. Fly condos and oviposition plates are soaked thoroughly with 10% Sparkleen (Fisher catalog number 04-320-4) solution and rinsed with distilled water after every use and allowed to air-dry (Kacsoh *et al.* 2015b, 10.7554/eLife.07423).

### RU486 feeding

We perform RU486 feedings in both assays used, as previously described (Kacsoh *et al.* 2015a, 1143-1157; Kacsoh *et al.* 2015b, 10.7554/eLife.07423). The method in which the RU486 is administered is similar in each behavior assay, but warrants two detailed methods. RU486 (Mifepristone) is used from Sigma (Lot number SLBG0210V).

For the ethanol seeking memory assay, we apply RU486 or vehicle only directly into the media. The caps for the fly corrals are made in the exact same manner as described above, but instead of using distilled water, an RU486 solution is used. The RU486 solution is prepared by dissolving 3.575 mg of RU486 in 800μL methanol (Fisher Scientific, Lot number 141313). This solution is added to 15.2 mL of distilled water. The total solution (16 mL) is thoroughly mixed and pipetted onto control food caps. For caps containing ethanol, the total solution of RU486 is changed to 14.992 mL and then pipetted onto the instant food. After pipetting the RU486 solution, we pipette 1.008 mL of 95% ethanol directly onto the food. Fresh RU486 solutions are prepared daily for experiments (Kacsoh *et al.* 2015a, 1143-1157).

For the oviposition depression, teaching and social learning assays involving inducible knock-down, we prepare the fly condos by measuring 0.375 grams of flaky instant blue Drosophila medium (Fisher Scientific Catalog number S22315C) into each well of The Fly Condo™ plates. For all food treatments, a total liquid volume of 2250 μL is directly pipetted onto the instant food. For experiments with RU486, an RU486 solution is used that is prepared by dissolving 3.575 mg of RU486 in 800μL methanol (Fisher Scientific, Lot number 141313). This solution is added to 15.2 mL of distilled water. The total solution (16 mL) is thoroughly mixed. From this mixed solution, we pipette 2250 μL onto the instant food. For plates containing no RU486 (vehicle, methanol only) 800μL methanol is mixed with 15.2 mL of distilled water. The total solution (16 mL) is thoroughly mixed. From this mixed solution, we pipette 2250 μL onto the instant food. RU486 solutions are prepped daily for experiments (Kacsoh *et al.* 2015b, 10.7554/eLife.07423).

### Statistical analysis

Statistical tests for behavioral assays are performed in Microsoft Excel. We use Welch’s two-tailed t-tests for all egg count data. P-values are calculated for comparisons between paired treatment-group and unexposed (Kacsoh *et al.* 2015b, 10.7554/eLife.07423; Kacsoh *et al.* 2015a, 1143-1157).

